# Retinal degeneration in a mouse model of CRB1 disease rescued by the photoreceptor-specific *CRB1-B* isoform

**DOI:** 10.64898/2026.08.05.743134

**Authors:** Ekta M. Dembla, Juan C. Valdez-Lopez, Christopher Kozlowski, Charlotte Reedy, Lin Yu, Mayur Dembla, Daniel Sul, Ariane Pendragon, Ying Hao, Jeremy N. Kay

## Abstract

Many genes involved in inherited diseases produce alternate mRNA isoforms that remain poorly characterized. Functional assessment of these isoforms could therefore unlock new insights into disease pathobiology or treatment. Here we investigated the function of the newly discovered “B” isoform of *CRB1*, a gene implicated in inherited retinal degenerations. *CRB1-B* is the most abundant retinal isoform, and differs from the canonical *CRB1-A* isoform in several important ways. Crucially, *CRB1-B* is the only isoform expressed by photoreceptors – the cells vulnerable to degeneration. Using a mouse model of CRB1-associated degeneration with degraded visual acuity, we find that restoring CRB1-B to photoreceptors prevents degeneration and completely rescues visual function, despite the absence of other isoforms. The therapeutic mechanism involves preservation of adherens junctions between photoreceptors and supporting glia: We establish that progressive loss of these junctions is a key pathobiological mechanism underlying photoreceptor death, and that junction loss is prevented by restoring photoreceptor expression of *CRB1-B*. Our findings nominate *CRB1-B* replacement as a promising gene therapy strategy. More broadly, they suggest that exploring disease gene isoform diversity offers an untapped opportunity to devise novel therapeutics.

## INTRODUCTION

Inherited retinal degenerations (IRDs) are a family of genetic disorders in which the light-sensitive photoreceptor cells of the retina are progressively lost, leading to profound visual impairment. IRDs are largely caused by mutations in single genes; over 300 such disease genes have been identified to date (1). For the vast majority of IRD-associated genes it remains unclear how disease-causing mutations alter specific biological processes leading to degeneration (2). In this way, IRDs are much like many other genetic disorders of the central nervous system: In general, far too little is known about the pathobiology of such diseases, hindering efforts to find cures. One possible reason for slow progress in determining the links between altered gene output and neurodegeneration is that the protein-coding mRNA output from many genes remains surprisingly poorly characterized. Defining the complete portfolio of key protein-coding mRNA isoforms encoded by disease genes is an essential step towards understanding how specific mutations cause disease.

Recently, our lab used long-read sequencing to profile the complete set of mRNA isoforms encoded by the IRD-associated gene *CRB1* (3). Mutations in human *CRB1* cause a broad spectrum of autosomal recessive clinical phenotypes, ranging from severe, early-childhood onset retinal degeneration (SECORD) or Leber congenital amaurosis (LCA) to milder forms such as retinitis pigmentosa (RP) or cone-rod dystrophy (4, 5). *CRB1* mutations are among the top ten most common causes of autosomal recessive IRDs (6). Efforts to understand the pathobiology of this disease have focused on the canonical mRNA isoform, *CRB1-A*, which encodes a transmembrane protein homologous to Drosophila Crumbs. Both Crumbs and its mammalian homologs have been shown to control apical epithelial polarity, with a particularly important role in assembly of apical junctions between epithelial cells (7, 8). The Crumbs intracellular domain, which is remarkably similar to that of human CRB1-A, is essential for these apical junctional functions. Accordingly, CRB1-A has been shown to traffic selectively to the apical membranes of Müller glia, a radial astroglial cell type that spans the entire retina(9, 10). Moreover, within this apical domain, CRB1-A localizes to the site of adherens junctions between Müller and photoreceptor cells that comprise the outer limiting membrane (OLM; Fig. 1A,B). These findings led to a model of disease pathogenesis whereby CRB1 is required for formation and/or maintenance of OLM junctions. However, this model remains unproven, as direct OLM damage in human patients is difficult to distinguish from indirect OLM damage caused by photoreceptor cell loss. Thus, animal models will be key to testing the idea that loss of CRB1 function causes pathogenic junctional phenotypes at the OLM. Unfortunately, past studies attempting to model this disease using *Crb1* mutant mice have not agreed on the extent to which *Crb1* is needed for OLM integrity or photoreceptor survival (10, 11).

**Figure 1.**
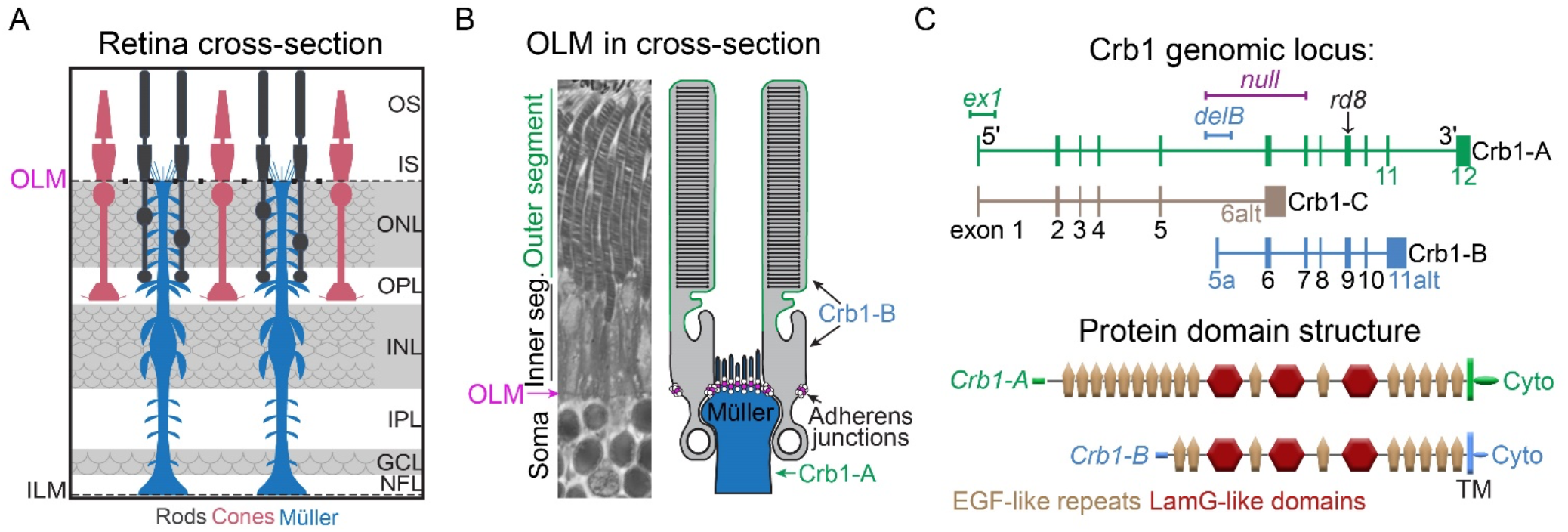
Retinal outer limiting membrane and the Crb1 gene. (A) Cross-section schematic of mouse retina in depicting cell types involved in formation of OLM: Rod and cone photoreceptors (gray, red), and Müller glia (blue). (B) Left: Thin plastic retinal section cropped to highlight OLM region. Right: Illustration of OLM anatomy and cell type composition, aligned to the section at left. Müller-photoreceptor junctions (magenta) surround each inner segment at OLM (magenta arrow). Cell types expressing each major CRB1 isoform are indicated. (C) Top, schematic of *Crb1* gene locus highlighting major isoforms (A, B, C). Vertical bars, exons. Horizontal bars, genetic lesions associated with mouse mutant alleles. Arrow, site of *rd8* point mutation that affects only *Crb1-A* and *Crb1-B* isoforms. The *null* mutation affects all isoforms. Bottom, domain structure of CRB1-A and CRB1-B proteins. Colors denote isoform-specific protein regions, including intracellular domains.

In our earlier study profiling *CRB1* isoforms in mouse and human, we identified a new isoform, *CRB1-B*, that is by far the most abundantly expressed retinal isoform in both species (3). CRB1-B has several key differences from the canonical CRB1-A isoform, including: 1) It lacks the conserved Drosophila intracellular domain mediating apical polarity functions; and 2) it is expressed by rod and cone photoreceptors, unlike CRB1-A which is selectively expressed by Müller glia. These differences imply that the new CRB1-B isoform has distinct biochemical and cell-type-specific functions. Disease-causing mutations are especially common within regions of the CRB1 protein shared by both the A and B isoforms, implying that both isoforms are involved in disease pathogenesis and that both could potentially be used in a gene therapy (5, 12). However, only CRB1-B is small enough to easily fit within an adeno-associated virus (AAV) gene therapy vector; and furthermore, AAV-mediated delivery of CRB1-A to mouse retina did not prove beneficial (13). Thus, delineating isoform-specific functions for CRB1-B will likely be highly relevant for understanding disease mechanisms and how to design gene therapy approaches. To date, however, CRB1-B functions within photoreceptor cells, and how these functions may be relevant to degeneration, have not been determined.

Here we sought to investigate the contribution of the CRB1-B isoform to disease pathology and photoreceptor degeneration. To do so, we needed to develop an improved mouse model of IRDs caused by CRB1 mutations. Two mouse *Crb1* mutant alleles have been used extensively in the literature (10, 11): 1) *Crb1^rd8^* (here denoted *rd8*), a spontaneously arising point mutation, which is predicted to affect both *Crb1-A* and *-B* but not other isoforms that are expressed in mouse retina (Fig. 1C); and 2) a targeted mutation of *Crb1* exon 1 (*ex1* allele), which was initially published as a knockout allele before identification of the *Crb1-B* isoform revealed that there are key isoforms that are not exon 1-dependent. Mutant mice homozygous for the *rd8* or *ex1* alleles do not have severe retinal degenerative phenotypes mimicking human patients. The *rd8* phenotype is more severe than *ex1*, in accordance with its location affecting both major isoforms (Fig. 1C, but even in these mice photoreceptor loss takes 1 year or more and is highly strain dependent (10, 11, 14). With this timeframe, experiments probing therapies or treatments are not practical. Using our *Crb1* isoform map, we developed a new *Crb1* mutant allele, denoted *Crb1^null^*(or simply *null*), that is predicted to eliminate expression of all isoforms (Fig. 1C). These mice exhibit a ∼20% decline in photoreceptor numbers by 3.5 months of age, suggesting they could provide an improved disease model compared to *rd8* or *ex1* (3).

Here we show that *Crb1^null^* mice have reduced visual acuity and progressive vision loss, validating them as a mouse model to investigate the pathobiology of CRB1 disease. We develop an improved quantitative assay for OLM junction defects and use it to show that *Crb1^null^* mice develop holes in their OLM long before the onset of degeneration. Remarkably, restoring only the *Crb1-B* isoform to its native cell type, the photoreceptors, is sufficient to largely rescue OLM pathology and photoreceptor survival – and to completely rescue visual acuity defects. These findings demonstrate a central role for the CRB1-B isoform in the pathobiology of CRB1 disease and provide a viable gene therapy strategy for CRB1-associated IRDs.

## RESULTS

### Progressive visual deficits in *Crb1^null^* mice

We previously showed that *Crb1^null^* homozygous mutant mice exhibit a mild (∼20%) loss of rod photoreceptors at 3.5 months of age (3.5 m) (3). To validate these *null* mice as an improved CRB1 disease model, we began our study by asking two questions: 1) Is this extent of photoreceptor loss sufficient to impair vision? and 2) Is degeneration progressive, as seen in human CRB1 IRD patients?

To test for visual impairment, we measured the visual acuity of *null* mice and strain-matched *Crb1^+/+^* controls (*+/+*) using an optomotor behavior assay (15). Experiments were conducted at mesopic light levels to probe rod photoreceptor function. We found that visual acuity of 3.5 m *Crb1 null* animals was significantly decreased relative to +/+ controls (Fig. 2A). This effect at the population level was accompanied by substantial individual variability: some *null* mice showed severe visual response impairment, while others performed similarly to control mice. This behavioral variability was consistent with anatomical variability we previously observed in *null* mutant photoreceptor counts (3). To ascertain whether anatomical variability could account for the variability of visual behavior, we prepared thin plastic sections from *null* mice with high vs. low acuity scores. Animals that scored poorly in the visual acuity assay reliably exhibited severe photoreceptor loss, resembling the severely degenerated examples shown in our previous study, whereas photoreceptors were largely preserved in animals with normal acuity scores (Fig. 2B). These findings demonstrate that photoreceptor loss in 3.5 m *Crb1 null* mice is sufficient to impair visual performance.

**Figure 2.**
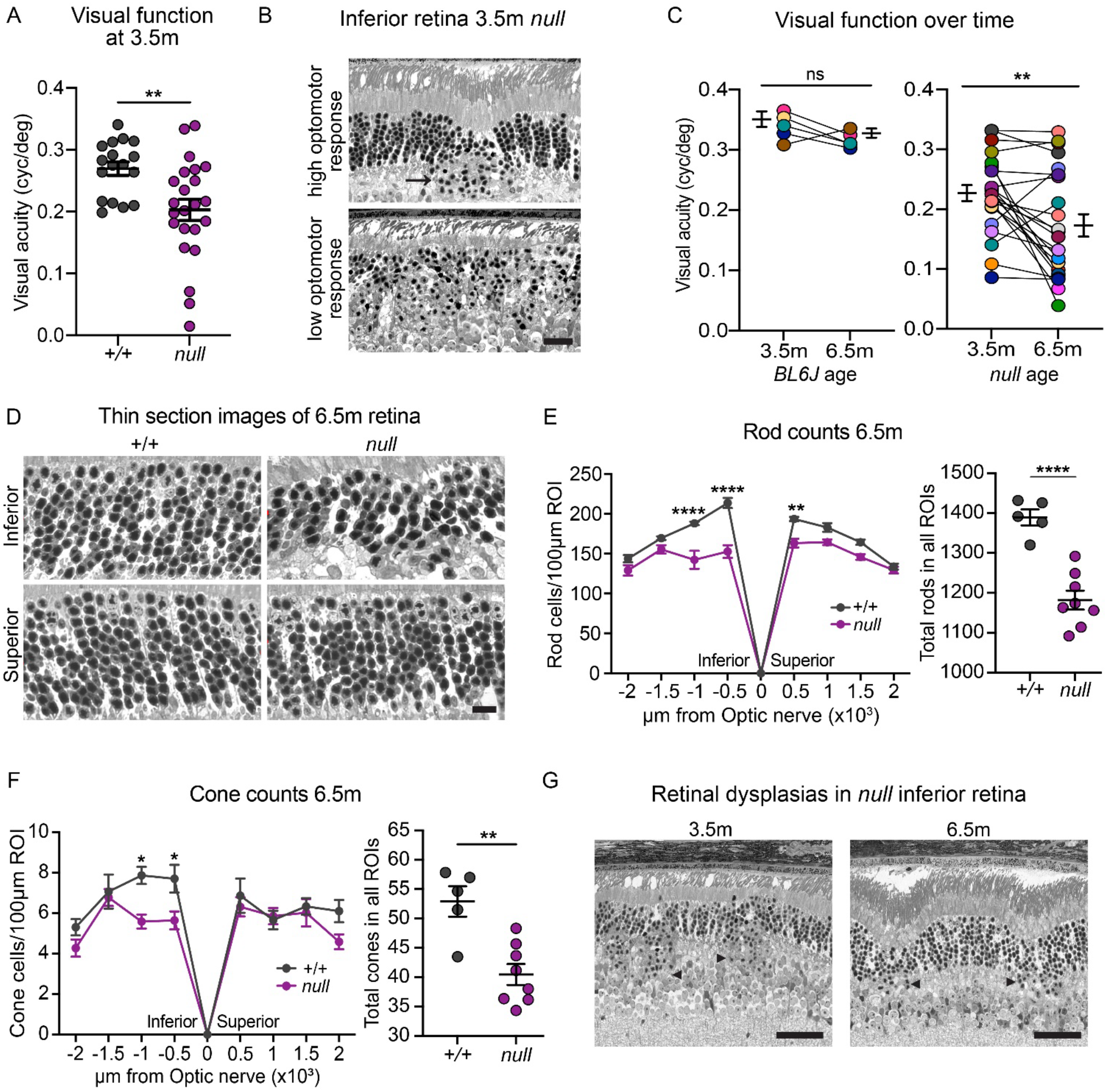
Retinal anatomy and visual behavior phenotypes in *Crb1^null^* mice. (A) Visual acuity thresholds measured using optomotor assay in *Crb1^null^* and strain matched *Crb1^+/+^* at 3.5 m. (B) ONL histology of 3.5 m *null* mutant mice that showed normal optomotor behavior (top) or impaired optomotor behavior (bottom). ONL thinning was well correlated with poor visual acuity, whereas hemi-rosette dysplasias (arrow) were often present in mice with normal acuity. (C) Comparison of visual acuity across two timepoints (3.5 m, 6.5 m) in cohorts of control C57Bl/6J (left) or *Crb1^null^* mice (right). Performance of individual mice (connected colored circles) and group averages are shown. Independent *null* cohorts were used (B, C). Statistics: (B) unpaired *t* test, ** *p* = 4.5×10^−3^; (C) paired *t* test, ns *p* (BL6J) = 1.6×10^−1^, ** *p* (*null*) = 3.6×10^−3^. (D-F) Degeneration of rods and cones in 6.5 m *null* mice. D: Representative thin plastic section images showing ONL of *null* or strain-matched +/+ controls. Images depict central retina, ∼500 µm inferior (top) or superior (bottom) to the optic nerve. E, F: Quantification of rod (E) or cone (F) numbers from regions of interest (ROIs) similar to D, sampling at regular intervals across the inferior-superior axis. Left, spider plots; right, total rod (E) or cone (F) nuclei counted across all ROIs. Degeneration in null retina is most pronounced in central retina, especially inferior to the optic nerve. Statistics: (E) Rod spider plot: two-way ANOVA with Bonferroni’s multiple comparisons test, ** *p* (at 500µm) = 0.004, **** *p* (at −500µm) <1.0×10^−7^, **** *p* (at −1000µm) = 3.3×10^−6^; Total rod count: unpaired *t* test, **** *p* =8.1×10^−5^; (F) Cone spider plot tests were the same as in E, * *p* (at −500 µm) = 0.036, * *p* (at −1000 µm) = 0.013; Total cone count: unpaired *t* test, *p* value ** = 0.002. (G) Representative images of hemi-rosette dysplasias (arrow heads) in *null* mutant retina at 3.5 m and 6.5 m. A greater tendency toward ONL laminar disruptions was noted at the latter age. Error bars: mean **±** S.E.M. Sample sizes: (B) n = 18 +/+, 23 *null*; (C) n = 5 BL6J, 23 *null*; (E-F) n = 5 +/+, 8 *null*. Scale bars: 10 μm (D); 50 μm (G); 20 μm (H, I).

To learn how visual function changes over time in *Crb1* mutants, we tested a separate cohort of control (C57Bl6/J) and *null* mice longitudinally, examining each animal at both 3.5 m and 6.5 m using the optomotor assay. To compare the extent of visual defects across *Crb1* alleles, this experiment also included *rd8* homozygotes. Whereas visual acuity did not change significantly across the 3.5 – 6.5 m interval in wild-type mice, acuity was significantly worse in *null* mutants at 6.5 m (Fig. 2C; Supplemental Fig. 1G). The majority of *null* animals performed worse at 6.5 m than 3.5 m (15/23), with some showing especially large declines in acuity (Fig. 2C). Other *null* mice performed similarly at the two ages; this reflected in part the fact that visual performance was already poor by 3.5 m for a subset of the *null* animals, diminishing resolution for detecting further declines (Fig. 2C). Unlike *null* mutants, *rd8* mutants did not exhibit loss of acuity between 3.5 m and 6.5 m; and moreover, *null* mutants exhibited worse acuity than *rd8* at each age, suggesting that the *null* mutation better captures the severity and progressive nature of human CRB1 disease (Fig. 2C; Supplemental Fig. 1G).

To bolster the behavioral finding that *null* mutant visual phenotypes are progressive, we next tested whether anatomical defects also worsen over time. As in our prior study (3), photoreceptor numbers were estimated in *null* mutants and +/+ controls by sampling at 450 µm intervals across thin plastic retinal sections. Human CRB1 disease involves loss of both rods and cones, but cones were not assessed in our last study; therefore, we first compared cone numbers across three timepoints: 0.5 m, 3.5 m, and 6.5 m. This analysis revealed that cones were unaffected in *null* mutants at the early timepoint and only mildly affected at 3.5 months: there was a trend towards cone depletion in inferior-central retina, but overall cone numbers did not differ from controls (Supplemental Fig. 1C,D). By contrast, in 6.5-month *null* mutants, the loss of inferior-central cones was far more severe and was accompanied by an overall decline in the cone population, suggesting that *null* mice undergo progressive cone loss (Fig. 2F).

We previously showed that rod numbers are decreased in *null* mice relative to +/+ by ∼18% on average, although there is substantial animal-to-animal variability (3). Here we extended this analysis by examining rod numbers at 0.5 m and 6.5 m. No difference in rod numbers were evident in 0.5 m mice, ruling out a developmental defect in photoreceptor production (Supplemental Fig. 1B). At 6.5 m, the rod degeneration phenotype was similar to 3.5 m: The average magnitude of rod loss relative to +/+ was ∼14%, but again we observed substantial animal-to-animal variability (Fig. 2E). As part of this variability, we noted an additional phenotype in 6.5 m *null* animals that could have obscured the extent of photoreceptor loss: A subset of eyes (n = 4/12) exhibited severe outer retinal defects, including warping and/or folding of the outer nuclear layer (ONL), which precluded the use of our sampling method and forced us to exclude them from photoreceptor quantification (Supplemental Fig. 1E). These dysplasias usually resembled, but were more severe than, the “hemi-rosette” dysplasias that are observed in 3.5 m *null* mutants (Fig. 2G) as well as other *Crb1* mutant alleles (10, 11, 16). However, in some cases the dysplasias were located in superior retina, which differs from the previously described hemi-rosettes which are exclusive to inferior retina (Supplemental Fig. 1E,F). While local dysplasias did not necessarily preclude inclusion of a given eye, the dysplasia phenotypes were far more severe, and affected far more of the OLM, in eyes deemed uncountable (Supplemental Fig. 1E). Thus, exclusion of these eyes likely introduced sampling bias leading to an underestimation of the extent of rod loss within our 6.5 m cohort. Altogether, therefore, the histological analysis suggests worsening degenerative phenotypes over time, supporting the behavioral studies (Fig. 2C). Together, these data indicate that *Crb1 null* mice recapitulate key aspects of the progressive rod-cone degenerative phenotype typical of human disease.

### Disruptions to the OLM precede degeneration in *Crb1^null^* mice

We next used the *Crb1 null* model to investigate candidate pathobiological mechanisms. We focused on OLM junctions because of the longstanding hypothesis that junctional defects are a key contributor to degeneration. OLM junctions localize to a specific domain of the photoreceptor inner segments, at the site where the inner segment contacts the apical domain of Müller the cells (Fig. 1B; Fig. 3A). These junctions can be labeled by immunostaining for OLM-selective junctional proteins, like ZO-1 and PALS1; or by using fluorescent probe-conjugated phalloidin, which strongly labels the OLM due to the enrichment of F-actin at adherens junctions (Fig. 3B, C; Supplemental Fig. 2A, B).

**Figure 3.**
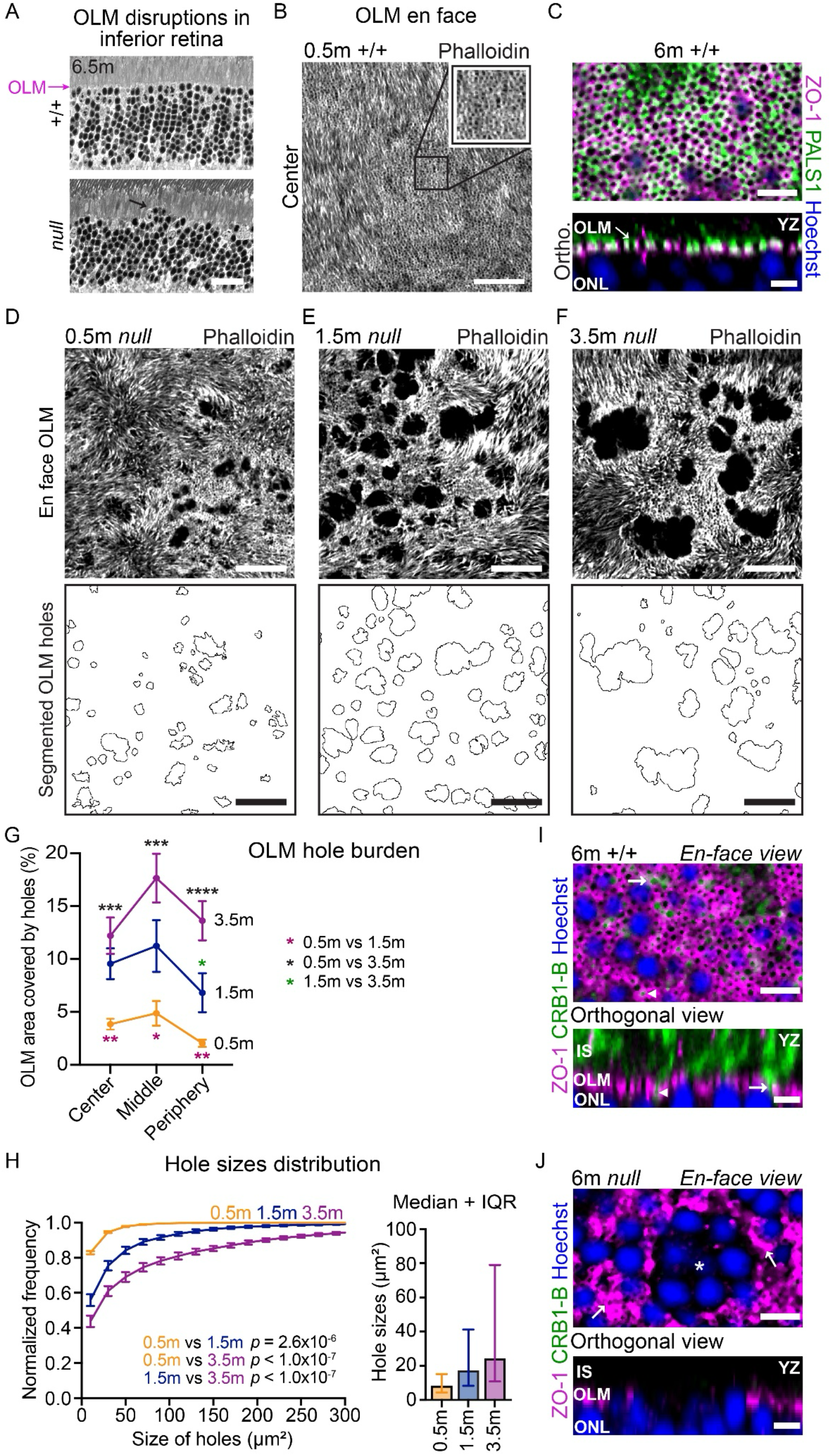
OLM pathology in Crb1 *null* mice. (A) Outer limiting membrane (OLM) histology in exemplary thin plastic sections from 6.5 m +/+ control (left) and *null* mutants (right). Note wavy appearance of mutant OLM and escape of nuclei into inner segment layer (arrow). (B) En face view of OLM junctions, labeled with phalloidin in retinal wholemount. Image is a single plane from a confocal Z stack acquired from P17 wild-type retina. Imaging plane is orthogonal to the illustration in A (red arrow in A indicates the approximate imaging plane). Inset, high-magnification view of boxed region. The OLM comprises a network of actin-rich rings surrounding inner segments. (C) En face confocal image of OLM in wild-type retina showing that rings express known OLM markers ZO-1 (magenta) and PALS1 (green). Bottom, orthogonal YZ projection of the confocal stack. PALS1-ZO-1 double-labeled rings (arrow) are positioned at apical surface of the ONL, where the OLM is located (A). Blue, Hoechst marking ONL nuclei. (D-F) OLM holes in *null* mutants. Top, representative en face images illustrating the *null* OLM phenotype at ages indicated. Each image is a single confocal slice at the OLM level, selected from a Z-stack acquired from central retina in phalloidin-stained retinal wholemounts. Bottom, segmentation and Z-projection of all OLM holes within the fields of view shown at top. Holes were never observed in wild-type (B; Supplemental Fig. 2A, 3A). (G) OLM hole burden analyzed over retinal eccentricities and over time. Hole burden was calculated from images similar to D-F, bottom row, as the percentage of the ROI area covered by holes. Note reduced hole burden in periphery at 0.5 m and 1.5 m. Overall hole burden increases over time. Statistics: One-way lognormal ANOVA with Šídák’s multiple comparisons test, 0.5 m vs 1.5 m – Center ** *p* = 0.002, Middle * *p* = 0.033, Periphery ** *p* = 0.004; 1.5 m vs 3.5 m – Periphery * *p* = 0.018; 0.5 m vs 3.5 m – Center *** *p* = 2.2×10^−3^, Middle *** *p* = 6.9×10^−4^, Periphery **** *p* = 1.4×10^−5^. (H) Distribution of hole sizes in Crb1 *null* retina over time represented in two ways. Left, cumulative distribution histogram (truncated at 300 µm). Right, Untruncated distribution represented as median ± IQR, whiskers – lower 25^th^ and higher 75^th^ percentile). Note increasing hole size over time. Statistics: Kolmogorov-Smirnov test for frequency distribution; see graph for *p* values. Measurements at all eccentricities were used for this analysis (for individual eccentricities see Supplemental Fig. 3C-E). (I, J) Expression of CRB1-B (green) at the OLM. I: En face and orthogonal views of ZO-1^+^ OLM junctions (magenta) in a 6m +/+ control. CRB1-B^+^ inner segments (green) protrude through OLM rings. Arrows of different thickness indicate the same inner segments viewed from each perspective. J: CRB1-B immunoreactivity is eliminated in *null* mice, and OLM holes lacking ZO-1^+^ junctions are evident (asterisk). Note also the clumped, abnormal appearance of ZO-1^+^ junctions adjacent to the OLM hole (arrows). Error bars: mean ± S.E.M (G, H, left); median ± IQR (H, right). Sample sizes: *null* 0.5 m n = 5; 1.5 m n = 7; 3.5 m n = 7. Scale bars: 20 µm (A, B, D-F); 15 µm (C, I, J en face); 10 µm (C, I, J orthogonal).

Using these markers, we first addressed CRB1 localization at the OLM. The CRB1-A isoform is well established as an OLM junctional protein (9); while we previously showed the B isoform localizes to inner segments, the presence of CRB1-B at the OLM has not been investigated. To do so, we raised a monoclonal antibody specific to the CRB1-B intracellular domain (Supplemental Fig. 2C). Immunohistochemistry on retinal wholemounts confirmed that CRB1-B is expressed by wild-type but not *null* photoreceptor inner segments, including at the sites where inner segments cross the OLM (Fig. 3I, J; Supplemental Fig. 2). Therefore, both isoforms are positioned to affect OLM junctional integrity.

We next assessed how loss of all CRB1 isoforms affects the OLM. Previous studies in *rd8* and *null* mutant mice used retinal cross-sections to show sporadic OLM defects – i.e., regions of the OLM in which photoreceptor-Müller adherens junctions are absent (3, 11). In these perturbed regions, ONL nuclei appear to “escape” into the ellipsoid zone through OLM holes. However, the extent of OLM damage was difficult to quantify from these cross-sectional images; and often the OLM phenotypes appeared quite subtle when junction defects are viewed in cross-section (Fig. 3A). To gain a more comprehensive view of OLM damage, we developed a new wholemount assay for OLM junctional defects in which damaged regions can be viewed en face, in their entirety, using phalloidin staining. En face imaging of wild-type or *Crb1^null/+^* heterozygotes stained in this manner revealed a ring-like pattern of phalloidin labeling at the OLM, with labeled rings surrounding each photoreceptor inner segment. (Supplemental Fig. 3B). These rings also express ZO-1 and PALS1, confirming their OLM identity (Fig. 3C). By contrast, in *null* mutants we observed striking OLM “holes” – defined as retinal patches in which inner segments lacked phalloidin-labeled junctions (Fig. 3D-F). ZO-1 labeling was also absent, supporting the conclusion that junctions are entirely missing within these lesions (Fig. 3I, J). Because this assay captures OLM holes in their entirety, we were able to quantify lesion severity by measuring 1) the number and size of OLM holes; and 2) overall OLM hole burden – i.e., the fraction of OLM territory in which junctions were missing.

Using this assay, we investigated whether OLM holes are present prior to onset of degeneration, as would be predicted if they are an underlying cause of photoreceptor death. OLM holes were quantified at postnatal day (P) 17 (also denoted as 0.5 m) – a developmental timepoint at which photoreceptor development is nearing completion. Even at this early age, OLM holes were already present in *null* mutants (Fig. 3D). By contrast, no holes were observed in +/+ control animals (Fig. 3B), nor in *Crb1^+/–^* heterozygotes (Supplemental Fig. 3A, B). We next tested whether the *null* OLM phenotype worsens overt ime, as might be expected if OLM defects contribute to progressive vision loss. By 1.5 m of age, OLM holes were larger and covered a larger proportion of retinal area than at 0.5 m, and a similar increase was again seen between 1.5 m and 3.5 m (Fig. 3D-H). The biggest change between 1.5 m and 3.5 m was in peripheral retina, which was relatively spared from OLM damage at 1.5 m but not 3.5 m (Fig. 3G; Supplemental Fig. 3C, D). These findings indicate that *Crb1* is required both for establishment of the OLM by 0.5 m, and for maintenance of OLM junctions in mature retina.

### *Crb1* OLM defects correlate with severity of degeneration

The presence of OLM holes prior to photoreceptor degeneration suggests that they could be an underlying cause of degeneration. If so, retinas with more severe OLM damage should ultimately undergo more degeneration. To test this prediction, we took advantage of the existence of multiple different *Crb1* mutant strains that vary in their extent of photoreceptor loss. If OLM holes are a cause of photoreceptor death, holes should be more severe in *null* mutants than in other *Crb1* mutant strains, such as *rd8* and the B-isoform-specific *Crb1^ΔB^* mutant (*ΔB*), that do not lose photoreceptors by 3.5 m (3).

To compare the severity of OLM defects across *Crb1* mutant strains, OLM anatomy was assessed early in development, at P17. This early timepoint was chosen to isolate direct effects on OLM anatomy from indirect effects arising from ongoing photoreceptor loss. Consistent with the idea that OLM hole severity is linked to degeneration, we observed that OLM defects were more severe in *null* mutants than in either of the other two strains. In *rd8* mutants, holes were present at P17 but they were significantly smaller than in *null* retinas (Fig. 4A-C; Supplemental Fig. 4A, B), with a strong trend toward fewer holes and lower OLM hole burden (Fig. 4D). The hole size difference was most pronounced in central retina, where *null* mutant holes were largest, but significant differences were detected across eccentricities (Supplemental Fig. 4A, B). Homozygous *ΔB* mutants, which selectively lack the *Crb1-B* isoform, had a relatively normal OLM: while some holes were observed, they were quite small (Fig. 4C; Supplemental Fig. 4C) and did not meaningfully alter OLM coverage (Fig. 4D). The *ΔB* OLM phenotype could be enhanced by additionally removing one copy of *Crb1-A* (*null/ΔB* mice, with 1 copy of *A* and 0 copies of *B*), but coverage was still minimally affected relative to *rd8* or *null* (Fig. 4D). Together, these findings indicate that the severity of degeneration and the severity of early OLM defects are correlated across mutant strains. Such correlations are consistent with the notion that OLM defects are an underlying cause of degeneration.

**Figure 4.**
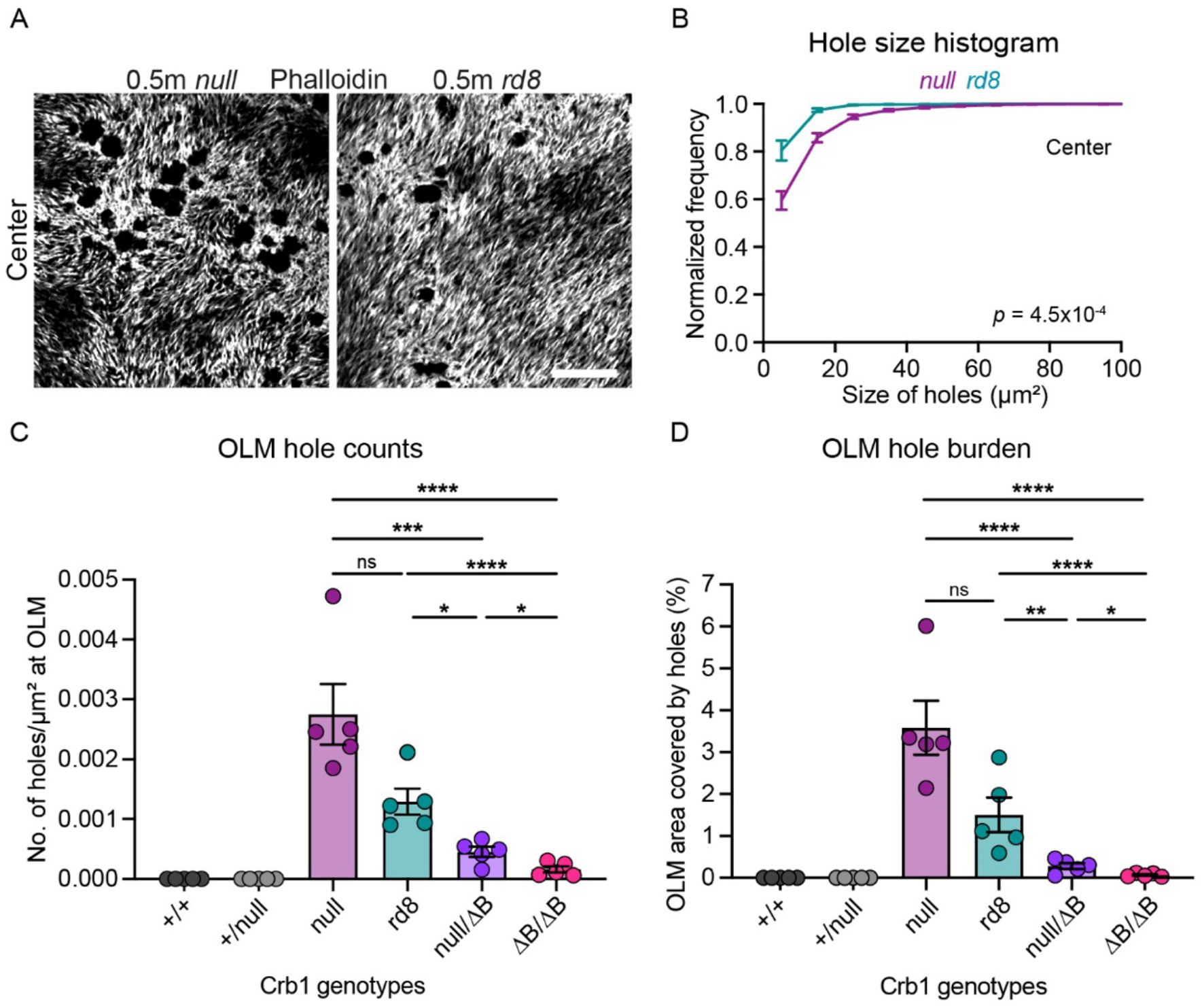
Comparison of OLM defects across *Crb1* alleles. (A) Representative en face views of OLM holes in retinal wholemounts from 0.5 m *Crb1^null^* and *Crb1^rd8^* mice. (B) Cumulative distribution histogram of hole sizes (truncated at 100 µm) in central retina of 0.5 m *Crb1^null^* and *Crb1^rd8^* mice, calculated from images similar to A. Statistics: Kolmogorov-Smirnov test for frequency distribution, for *p* value see graph. (C) OLM hole frequency in 0.5 m mice with different *Crb1* allelic combinations, calculated as holes/μm^2^ from ROIs similar to A. No holes were observed in wild-type or +/*null*. Statistics: One-way lognormal ANOVA with Šídák’s multiple comparisons test, *null* vs *rd8* – ns *p* = 0.212, *null* vs null/ΔB *** *p* = 2.6×10^−4^, *null* vs ΔB/ΔB **** *p* = 7.0×10^−7^; *rd8* vs null/ΔB * *p* = 0.026; *rd8* vs ΔB/ΔB **** *p* = 2.7×10^−5^; and null/ΔB vs ΔB/ΔB * *p* = 0.019. (D) OLM hole burden across *Crb1* allelic combinations, calculated as hole area/ROI area. Statistics: One-way lognormal ANOVA with Šídák’s multiple comparisons test, *null* vs *rd8* – ns *p* = 0.254, *null* vs null/ΔB **** *p* = 5.1×10^−5^, *null* vs ΔB/ΔB **** *p* = 3.0×10^−7^; *rd8* vs null/ΔB – ** *p* = 0.003; *rd8* vs ΔB/ΔB – **** *p* = 7.0×10^−6^; and null/ΔB vs ΔB/ΔB – * *p* = 0.028. Error bars: mean ± S.E.M. Sample size: n = 5 *null*; n = 5 *rd8*. Scale bar: 20 μm.

### OLM pathology is accompanied by microglia recruitment

The presence of OLM defects in 1.5 m *null* mice suggest that photoreceptor pathology may already be occurring at this time, despite the fact that we did not detect cell loss until 3.5 m. To investigate this possibility, we examined the distribution of retinal microglia. Prior studies have demonstrated that one key hallmark of photoreceptor damage is the recruitment of microglia to the outer retina: Upon photoreceptor injury, microglia leave their homeostatic niche within the outer plexiform layer (OPL) and migrate across the ONL to take up residence at sites of tissue damage within the subretinal space (17–19). To learn whether this phenomenon also occurs in *Crb1^null^* mice, we examined the laminar distribution of microglia in retinal cross-sections from 1.5 m *null* mutants. In line with other degeneration models, Iba1^+^ microglia were observed in the *null* outer retina – not only within the ONL but also crossing into the subretinal space (Fig. 5A). These Iba1^+^ cells had a migratory morphology and expressed CD68, a marker of migrating microglia (19), suggesting that they were not macrophages but instead were microglia recruited into the outer retina from the OPL (Fig. 5A). Some Iba1^+^ cells within the ONL had phagocytic cups surrounding photoreceptor nuclei, indicating that they were responding to tissue damage by clearing damaged cells (Fig. 5A, yellow arrow). By contrast, strain matched controls never showed any Iba1^+^ cells in outer retina; they were confined to the OPL. These findings suggest that *null* outer retina is already damaged by 1.5 m of age.

**Figure 5.**
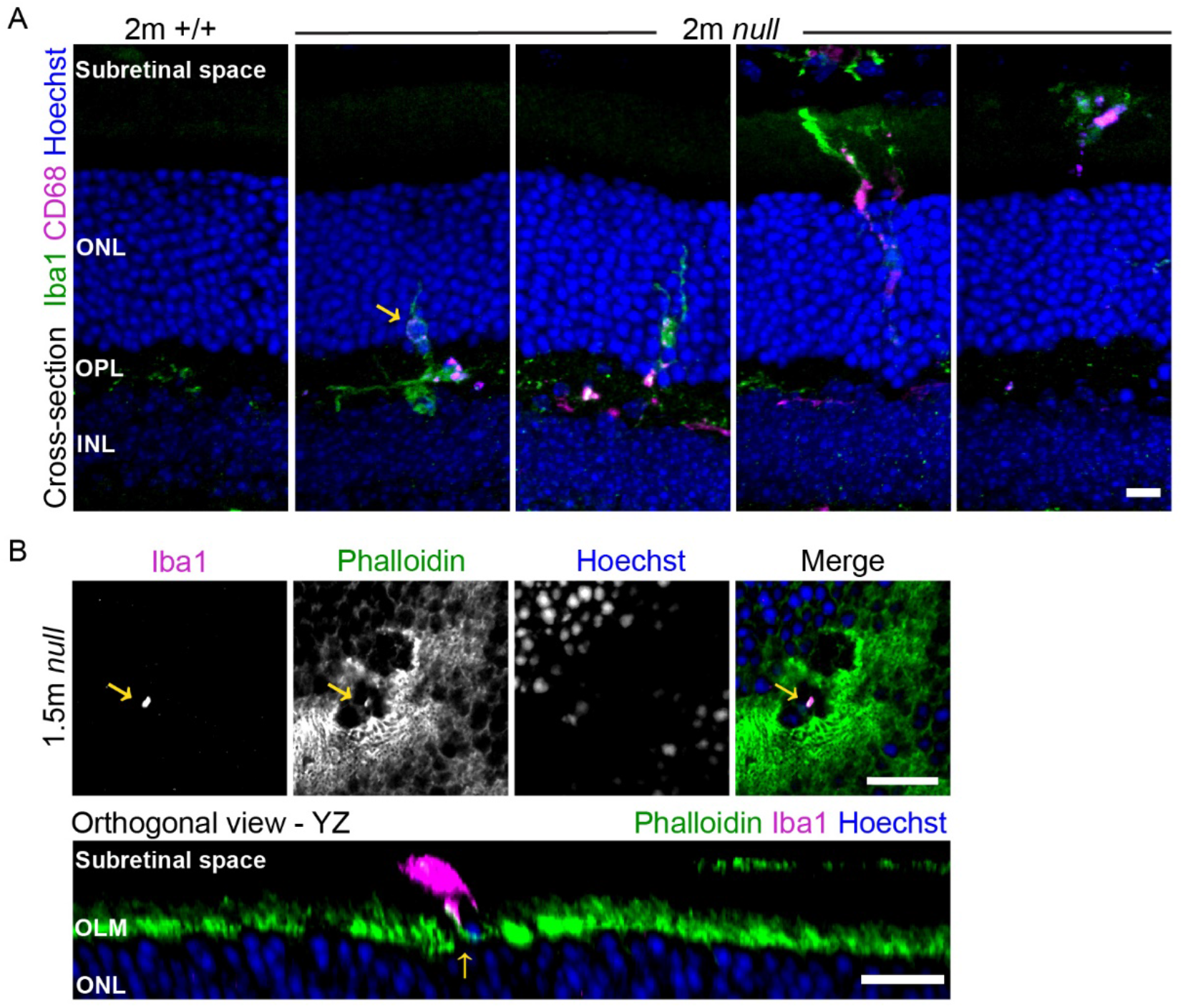
Microglia associate with OLM holes in *Crb1^null^* mice. (A) Cross-sections from 2m. *Crb1* +/+ or *null* retinas stained for microglia markers Iba1 (pan-microglia) and CD68 (lysosomes of phagocytic microglia). Wild-type microglia are confined to OPL, whereas in *null* mutants, microglia move through ONL into the subretinal space. Arrow, phagocytic cup engulfing ONL nucleus. (B) Top: En face view of a representative OLM hole from a 1.5 m *null* wholemount labelled with phalloidin and Iba1. Panels show the same field of view from a single confocal slice. Bottom: orthogonal YZ projection of the same OLM hole. An Iba1+ cell is located directly above the hole in the subretinal space, and projects processes into the hole (arrow). Scale bars: 50 μm (A); 20 μm (B, top), 10 μm (B, bottom).

We next addressed the possibility that microglia recruitment occurs selectively at sites of OLM damage, as would be predicted if OLM holes are indeed a proximal cause of degeneration. To test this idea, we examined the distribution of Iba1^+^ cells relative to OLM holes in *Crb1^null^*retinal wholemounts. At 1.5 m, we did not observe a large number of outer retinal Iba1^+^ cells – typically 5 or fewer that remained with the retina during the wholemount preparation. This count could underestimate the true number, because some microglia that had exited to the subretinal space may have been lost during the wholemount preparation, as the retina was removed from the eyecup. However, these few outer retinal Iba1^+^ cells were reliably localized to OLM holes (Fig. 5B; n = 34 microglia from 5 mice, 27 with cell bodies aligned to holes, 7 with processes extending through holes). Three-dimensional reconstructions of confocal stacks showed that most of these cells were polarized along the vitreal-scleral axis, with processes extending across the OLM(Fig. 5B). This morphology suggests that the Iba1+ cells were in the process of migrating from the OPL towards the outer retina, crossing the OLM at the hole sites. We did not observe Iba1^+^ cells crossing the OLM at sites without holes, suggesting that the microglia behavior was specific to regions with junctional defects. In some animals with particularly large OLM holes, Iba1^+^ cells ramified arbors within the holes, filling them (Supplemental Fig. 5). Altogether, these observations support the idea that microglia are responding to tissue damage caused by the presence of OLM junction defects.

### Transgenic strategy to restore CRB1-B in photoreceptors

We next leveraged our *Crb1^null^* disease model to investigate the function of the photoreceptor-specific CRB1-B isoform. Because *null* mutants have a stronger degeneration phenotype than the other *Crb1* alleles in which various isoforms remain intact, it seems likely that the CRB1-B isoform works in concert with other isoforms to promote photoreceptor survival. If this is the case, restoring either the A or the B isoform could be a viable therapeutic strategy. Unfortunately, efforts to restore CRB1-A into *Crb1 ex1* mutant mice were so far not successful, perhaps due to the size of the CRB1-A cDNA which does not easily fit into AAV vectors (13). The smaller CRB1-B isoform, by contrast, is not expected to have that limitation. We therefore focused on the possibility that restoring CRB1-B into its native cell type could ameliorate retinal phenotypes of *null* mutant mice.

To test this idea, we generated transgenic mice in which the well-characterized *Rhodopsin* promoter (20) was used to drive expression of a *Crb1B-IRES-GFP* construct selectively within rod photoreceptors (Fig. 6A). We term this transgenic construct Rho-Crb1B. The *IRES-GFP* portion was included to facilitate screening of transgenic founder lines, without altering the sequence of the CRB1-B protein itself. We obtained 12 founder lines which were screened at P21 for CRB1-B and GFP expression, using Western blot and immunohistochemistry (Fig. 6A; Supplemental Figs. 2C, 6D). Two lines were identified in which GFP, and CRB1-B were selectively expressed by the vast majority of rod photoreceptors, without off-target expression in other retinal layers (Fig. 6A; Supplemental Fig. 6D) or tissues (Supplemental Fig. 6E). Because the lines were similar, we chose one of them (line 9) for use in all downstream studies. Western blots demonstrated that CRB1-B expression was restored in *null;* Rho-CRB1B animals (here denoted *B-rescue* mice) to a higher level than in wild-type animals (Fig. 6C). However, the presence of excess CRB1-B protein is not detrimental to photoreceptor survival or morphology, as shown by comparison of transgene-positive and transgene-negative *Crb1^+/+^* mice (Supplemental Fig. 6A-C).

**Figure 6.**
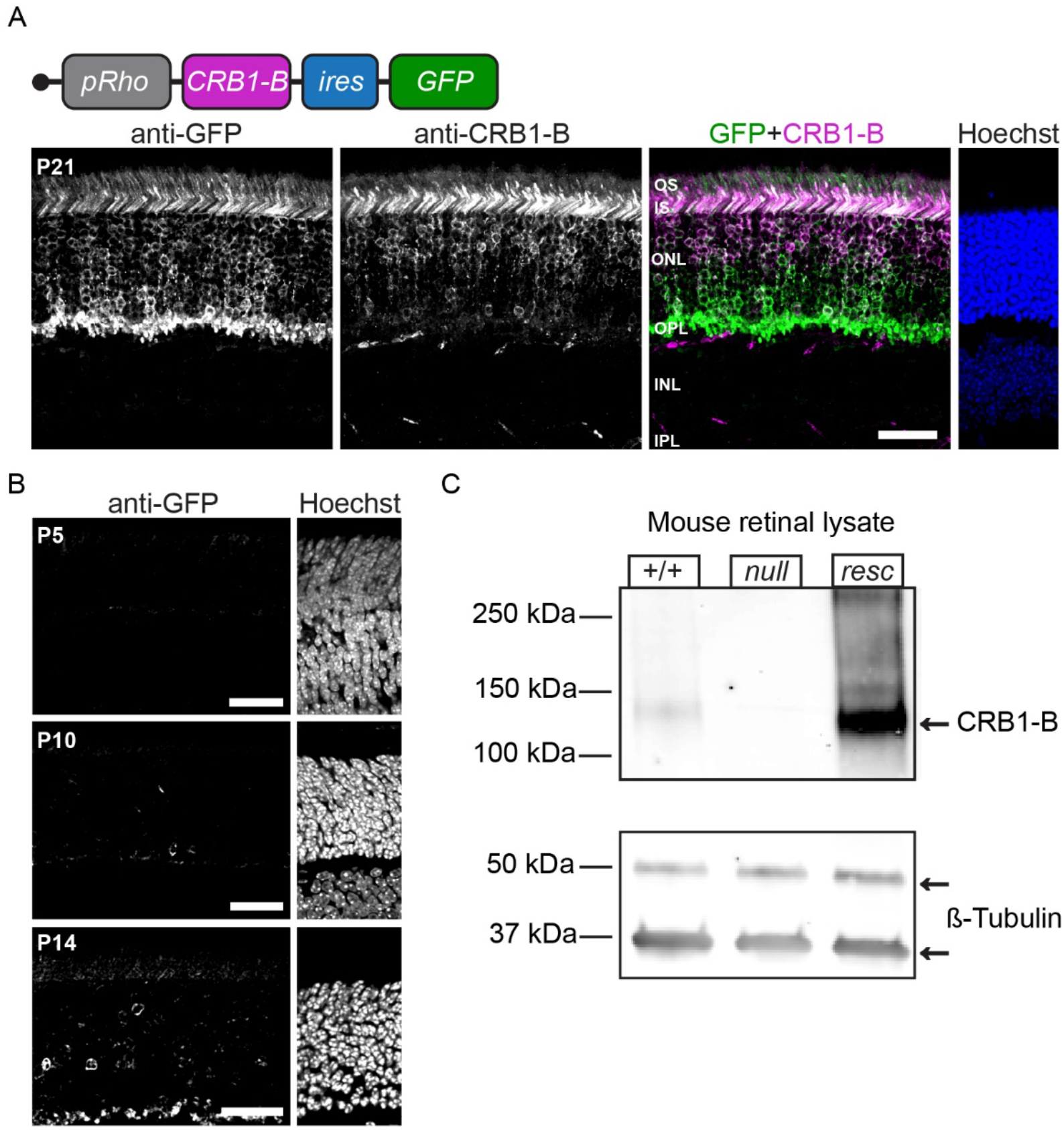
Rho-Crb1B transgenic rescue mice. (A) Top: transgenic rescue construct design. Bottom, vibratome section of P21 transgenic retina (*Crb1^+/+^*; Rho-CRB1B) stained for GFP (green) and CRB1-B (magenta). Each panel shows same field of view. Transgene is selectively expressed by rods; somata in ONL and inner/outer segments (IS/OS) are immunoreactive for both GFP and CRB1-B. (B) Timing of transgene onset, illustrated in vibratome sections from transgenic retina (*Crb1^+/+^*; Rho-CRB1B) at specified ages. Early expression is evident by P14 when small number of rods express GFP until a high expression at P21 (A). Images are representative of 2 mice at each time point. (C) Western blot of retinal lysates from Crb1 +/+, *null*, and *B-rescue* (*resc*) mice (i.e., *null* mice carrying Rho-Crb1B transgene). Blot was probed with antibodies to CRB1-B and β-tubulin (images depict the same membrane, see full blot in supplemental material). CRB1-B is absent from *null* retina but restored to greater than wild-type levels in *B-rescue* mice. Scale bars: 30 μm.

To determine the timing of rescue, we next characterized the onset of transgene expression. At P5 and P10, GFP was not yet detectable, consistent with prior reports as to the onset of Rhodopsin protein expression (21). By P14, a small number of rods expressing GFP were observed; and by P21, most rods had initiated GFP expression (Fig. 6A, B). These findings indicate that the Rho-Crb1B transgenic line restores CRB1-B protein expression to *null* mutants during the third postnatal week, at least several days after the OLM starts to form (P7; (22)).

### Restoring CRB1-B to rods ameliorates degeneration and rescues visual acuity

To learn whether the Rho-CRB1B transgene can influence photoreceptor survival, we generated cohorts of sibling *null* and *B-rescue* mice, and evaluated their retinal phenotypes at 3.5 m. As a control cohort, we also generated *Crb1^+/+^* mice that were half-siblings to the *null* and *B-rescue* groups. Photoreceptor counts in these three cohorts confirmed our prior results (3) (Fig. 2): The number of rods was significantly lower in 3.5 m *null* mutants relative to wild-type controls, with the largest effects in central retina, especially on the inferior side (Fig. 7A,B). Comparing these two groups to *B-rescue* mice, we found that reintroduction of CRB1-B led to a significant increase in rod numbers (Fig. 7B). While the numbers did not return all the way to wild-type levels, there was an improvement in photoreceptor survival across retinal eccentricities (Fig. 7B). Furthermore, we also noted a decrease in the frequency of rosette-like dysplasias, indicating that multiple aspects of the *null* mutant anatomical phenotype could be rescued by restoring CRB1-B (Fig. 7C,D).

**Figure 7.**
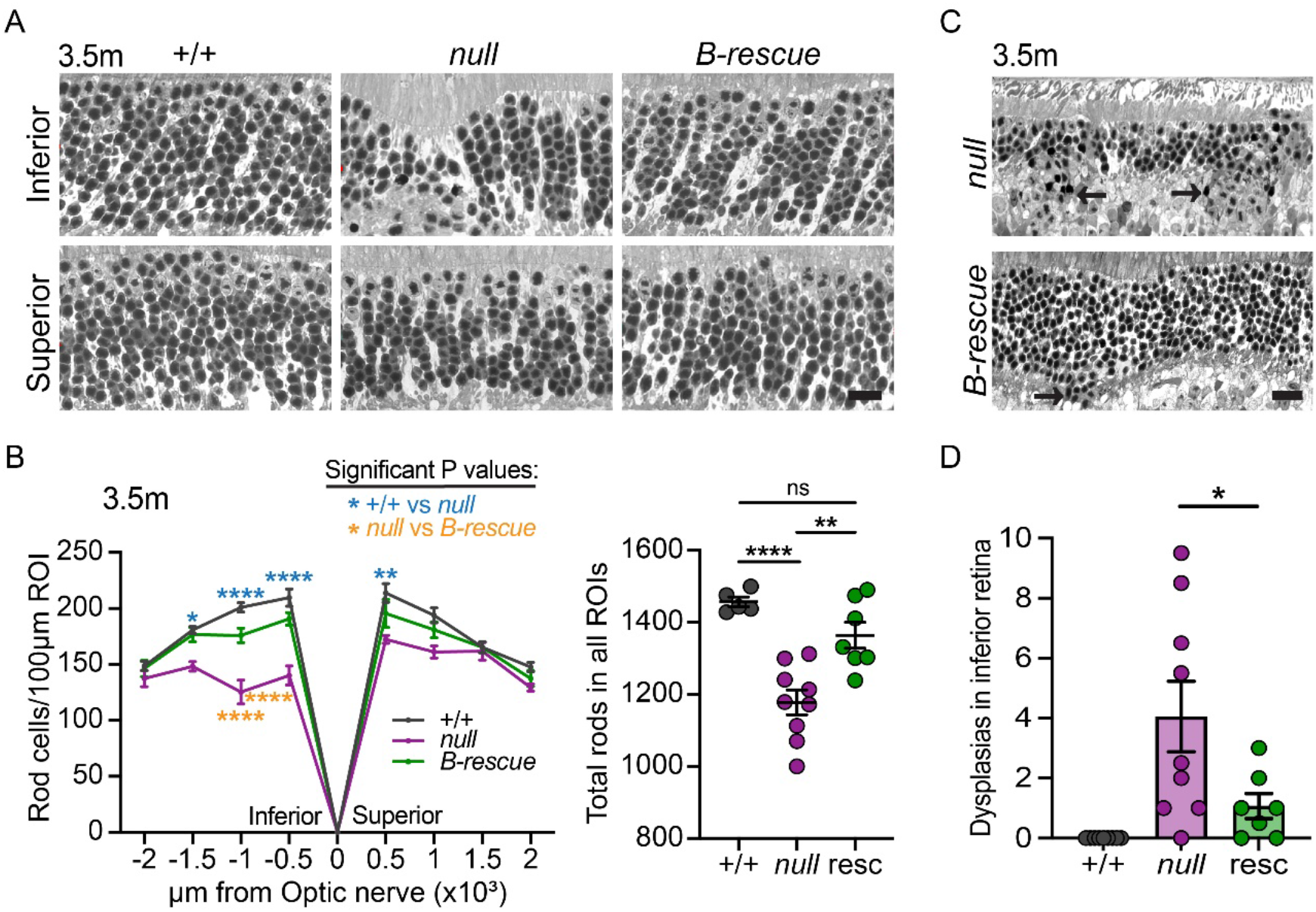
Restoring CRB1-B to rods ameliorates degeneration. (A) Representative images showing ONL of +/+, *null* and *B-rescue* mice at 3.5 m. Images depict central retina, inferior (top) or superior (bottom) to the optic nerve. (B) Quantification of rod numbers from images similar to A, sampling at regular intervals across the inferior-superior axis. Left, spider plots; right, total rod nuclei counted across all ROIs. Statistics: Spider plot: two-way ANOVA with Bonferroni’s multiple comparisons test, +/+ vs *null*, **** *p* (at −500 µm) = 2.0×10^−7^, **** *p* (at −1000 µm) <1.0×10^−7^, * *p* (at −1500 µm) = 0.031, ** (at 500 µm) = 0.008; *null* vs *resc*, **** *p* (at −500 µm) = 4.3×10^−5^, **** *p* (at −1000 µm) = 9.9×10^−6^. Total rod count: one-way ANOVA with Šídák’s multiple comparisons test, +/+ vs *null* **** *p* = 7.8×10^−5^, *null* vs *resc* ** *p* = 0.002 and ns *p* = 0.256. (C, D) Rescue transgene diminishes rosette dysplasia phenotype. C: Representative thin plastic section images of ONL hemi-rosettes from inferior retina of *null* or *B-rescue* mice. D: Quantification of hemi-rosette dysplasias in +/+, *null* and *B-rescue* mice. These dysplasias were not observed in superior retina. Dysplasias from inferior retina were counted from one retinal section/eye from one animal and averaged per animal. Statistics: unpaired t test, *null* vs *resc*, * *p* = 0.049. Error bars: mean **±** S.E.M. Sample sizes: (B, D) n = 5 +/+, 9 *null*, 7 resc. Scale bars: 10 µm (A), 20 µm (C).

To ask whether these anatomical improvements could be attributed to a rescue of OLM junctional defects, we next quantified OLM hole size and coverage in *null* and *B-rescue* mice. OLM holes were not completely eliminated in *B-rescue* animals (Fig. 8A; Supplemental Fig. 7), consistent with the fact that the OLM has already started to form by the time of transgene expression onset (Fig. 6A,B; (22)). However, holes were smaller in 1.5 m *B-rescue* animals than in *null* mutants, suggesting a positive effect on OLM junction maintenance (Fig. 8B). The positive effect was due largely to central and middle retinal eccentricities (Supplemental Fig. 7B), which are the most severely affected at this age (Fig. 2G). While the magnitude of the hole size difference was small at 1.5 m (Fig. 8B), effects of the transgene on OLM maintenance were much more striking when we compared 1.5 m and 3.5 m animals (Fig. 8B-E). Over this time, hole size and overall hole burden became notably worse in *null* mutants, but the presence of the *B-rescue* transgene entirely blocked this effect (Fig. 8E; Supplemental Figs. 7,8). Together, these findings demonstrate that restoring CRB1-B to rods in *Crb1^null^* mice prevents OLM damage and has a significant positive effect on photoreceptor survival.

**Figure 8.**
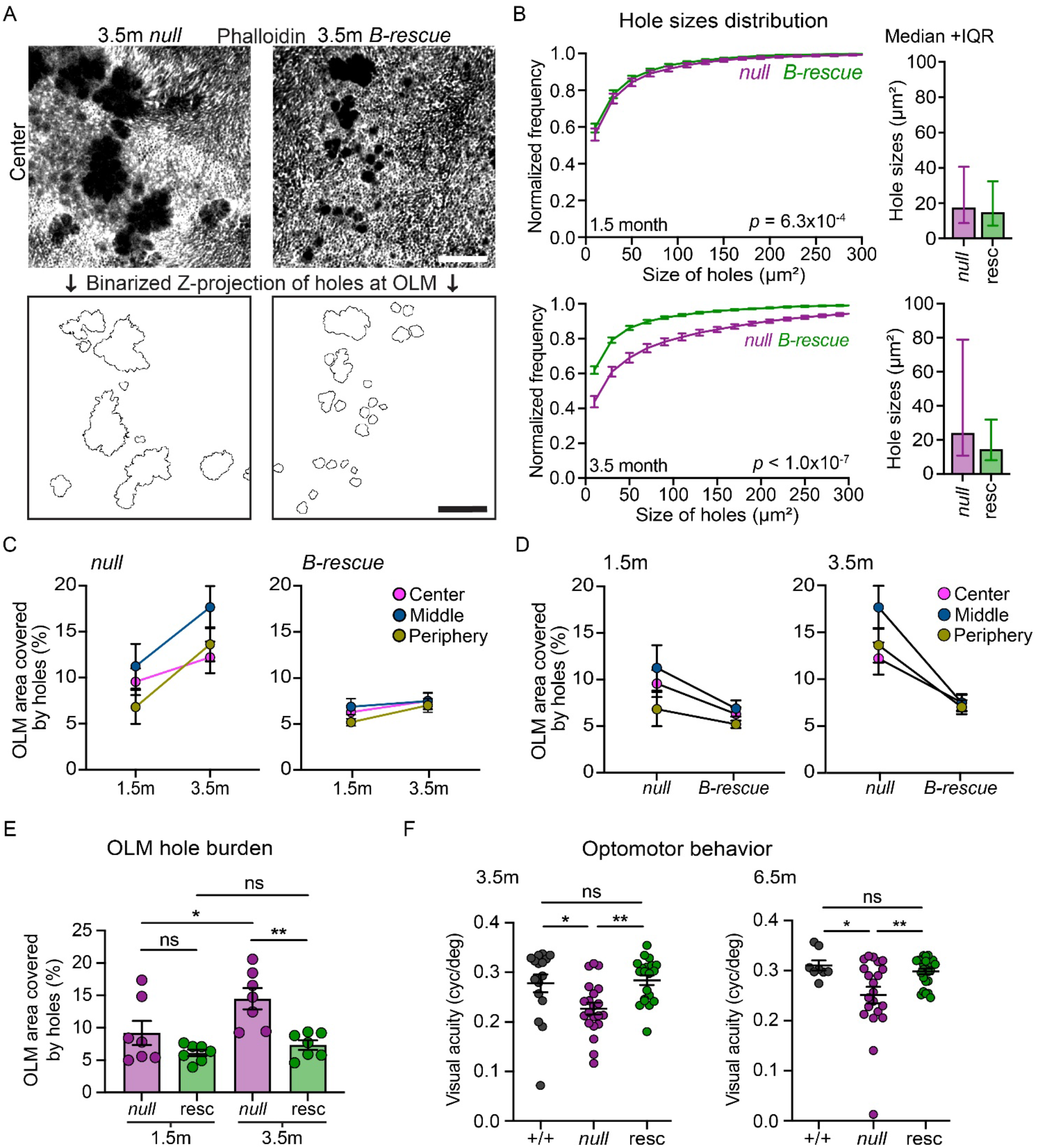
Restoring CRB1-B to rods arrests OLM junction loss and rescues visual acuity. (A) Representative images showing OLM hole phenotype in *null* and *B-rescue* wholemount retinas at 3.5 m. Top: En face phalloidin labeling in a single confocal slice, selected from a Z-stack encompassing the OLM. Bottom: The same fields of view showing segmented OLM holes Z-projected across the full stack. (B) Left, cumulative distribution histogram of hole sizes (truncated at 300 µm) in Crb1 *null* and *B-rescue* central retina at ages indicated. Null data are re-plotted from Fig. 3H. Hole sizes increase between 1.5 m and 3.5 m in *null* but not *B-rescue* mice. Right, hole sizes distribution represented as median **±** IQR (whiskers – lower 25^th^ and higher 75^th^ percentile). Statistics: Kolmogorov-Smirnov test for frequency distribution, for *p* values see graph. (C-E) Rescue transgene ameliorates OLM hole burden. C, D: Hole burden at 1.5 m and 3.5 m in *null* and *B-rescue* mice, plotted for each retinal eccentricity. C groups by genotype for comparison across age; D shows the same data grouped by age for comparison across genotype. E: Total OLM hole burden (pooled across eccentricities). Hole burden increases between 1.5 m and 3.5 m in *null* mice but remains stable in *B-rescue* mice. Null data are re-plotted from Fig. 3G and Supplemental Fig. 3D. Statistics: One-way lognormal ANOVA with Šídák’s multiple comparisons test, *null* vs *resc* (1.5 m), ns *p* = 0.338, 1.5 m vs 3.5 m (*null*), * *p* = 0.036, *null* vs *resc* (3.5 m), ** *p* = 0.005, and 1.5 m vs 3.5 m (*resc*), ns *p* = 0.832. (F) Rescue transgene preserves visual function. Visual acuity thresholds were measured from +/+, *null,* and *B-rescue* mice at 3.5 m (left) and 6.5 m (right) using optomotor assay. Separate cohorts of mice were assessed at each age. Statistics: One-way ANOVA with Šídák’s multiple comparisons test. 3.5 m: +/+ vs *null*, * *p* = 0.026 and *null* vs *resc*, ** *p* = 0.006. 6.5 m: +/+ vs *null*, * *p* = 0.024 and *null* vs *resc*, ** *p* = 0.009. Error bars: mean **±** S.E.M except in B as indicated (median **±** IQR). Sample sizes: (B-E) n = 7 *null*, 7 resc; (F) 3.5 m n = 16 +/+, 21 *null*, 20 resc; 6.5 m n = 8 +/+, 21 *null*, 25 resc Scale bars: 20 µm

Finally, we investigated whether the improvement in photoreceptor survival mediated by the CRB1-B rescue transgene was sufficient to rescue visual function. To this end, the cohorts of wild-type, *null*, and *B-rescue* mice were tested at 3.5 m for visual acuity using the optomotor assay. Remarkably, visual acuity was rescued entirely back to wild-type levels in *B-rescue* mice (Fig. 8F). Moreover, we found that this preservation of visual function was still present in a separate cohort of animals aged to 6.5 m (Fig. 8F). Together, these findings indicate that the rod-targeted CRB1-B rescue strategy not only ameliorates degenerative anatomical phenotypes in *null* mice but also suffices to provide long-lasting preservation of visual function.

## DISCUSSION

In this study we demonstrate the functional importance and therapeutic potential of the photoreceptor-specific CRB1-B isoform. When our lab first described this isoform 6 years ago (3), the prevailing model of CRB1 disease pathobiology was focused on the Müller glia-specific CRB1-A isoform (10, 16). It was known, based on the phenotype of *Crb1^rd8^* mice, that CRB1 was important for structural integrity of the OLM (11). But the connection between OLM damage and photoreceptor degeneration was difficult to establish – both because of the mild and inconsistent degenerative phenotype in *rd8* mice (11, 14, 23, 24), and because other *Crb1* mutant alleles lacked substantial OLM damage (9, 10, 16). Identification of the CRB1-B isoform allowed us to overcome this difficulty by generating *Crb1^null^* mutant mice, which we show here to be an improved model of CRB1 disease.

Using this new disease model, we show a strong correlation between the severity of OLM damage and the severity of photoreceptor loss. The link between OLM defects and degeneration was observed in three different scenarios: 1) Comparison across an allelic series of 5 different *Crb1* mouse mutants, which co-varied in their extent of OLM and photoreceptor defects. 2) Comparison across time and retinal eccentricity in *Crb1^null^* mutants: We observed that OLM holes grew larger with age and recruited microglia in a manner that preceded photoreceptor death. Further, we observed that central retinal regions, which degenerate first, are also the first to sustain substantial OLM damage. 3) Comparison between *null* and *Crb1B-rescue* transgenic mice, in which restoring expression of CRB1-B to rods caused an improvement both in the extent of OLM damage and photoreceptor survival. Together, these mouse findings strongly suggest that OLM damage is an underlying cause of degeneration, implying that this same pathobiology could underlie human CRB1 disease. Furthermore, our results suggest that restoring CRB1-B to photoreceptors is a viable strategy for treating CRB1 disease. These insights into a rare inherited retinal degeneration hold lessons for other genetic diseases: Our work illustrates how a better mechanistic understanding of such diseases, and new therapeutic strategies, can emerge from a more complete understanding of disease gene isoform diversity.

### The *Crb1^null^* model highlights the OLM as a key site of CRB1 disease pathology

A central finding of this study is that *Crb1^null^* mutant mice provide the first reliable animal model of human CRB1 disease. The major improvements compared to previous models are: 1) Earlier and more reliable photoreceptor degeneration; and 2) Functional deficits, measured through optomotor tests of visual acuity. Compared to *null* mice, *Crb1^ex1^*mutant mice have far less severe OLM junctional phenotypes, with minimal photoreceptor loss or degradation of visual function (10). *rd8* mice do exhibit OLM junctional defects, but we show here that these defects are not as severe as in *null* mice. Furthermore, photoreceptor loss in *rd8* takes far longer than in *null* mutants – a year or more – and is highly strain-dependent, with minimal degeneration seen on the C57Bl6 background (11, 23). By contrast, *null* degenerative phenotypes are consistent both on a mixed C57Bl6-SJL background (3) and after extensive backcrossing to C57Bl/6J (this study). Because degeneration in *rd8* mutants is so variable, few studies have endeavored to test visual function. Here we directly compared visual acuity in *rd8* and *null* mutants, and confirmed that functional deficits are also more severe in the *null*. Thus, both anatomical and functional measures show that *null* mutants have stronger disease-related phenotypes than *rd8* or *ex1* mutants.

It important to note that *Crb1^null^* animals do not model all aspects of the human disease: A large subset of CRB1 patients, with LCA or SECORD diagnoses, exhibit retinal laminar disorganization accompanied by a very early onset degeneration (5, 12, 25). With its later disease onset and preserved lamination, the *null* mice do provide a good model for CRB1 patients that have received RP or rod-cone dystrophy diagnoses.

Using the *null* mouse model, we demonstrate a role for formation and/or maintenance of OLM junctions in the pathobiology of CRB1 disease. This idea had been proposed previously, based on the localization of CRB1-A protein to the Müller side of the OLM junctions, but definitive tests of the idea were not possible until we developed a mouse model that shows reliable degeneration, as well as a quantitative assay for OLM damage. The data presented here are consistent with a model in which OLM junction integrity depends on the presence of different CRB1 isoforms on opposite sides of the OLM junction – *A* on the Müller cell side and *B* on the photoreceptor side. Loss of either isoform alone has minimal effects on junction integrity (Fig. 4; (10)), suggesting that each CRB1 isoform possesses the ability to regulate junction formation both within its own cell type and in a transcellular manner. Future studies will be needed to elucidate how CRB1 isoforms achieve these critical biochemical functions.

Our data strongly support the idea that sites lacking OLM junctions are subject to tissue damage leading to degeneration. However, other CRB1 pathobiological mechanisms are still possible. Prior studies have emphasized that loss of CRB1 leads to hemi-rosette dysplasias (10, 11). These are unlikely to be a major independent cause of degeneration, given that they are prevalent even in the phenotypically mild *ex1* mutant; however, our rescue studies demonstrated that the incidence of these dysplasias was also reduced in *B-rescue* mice, similar to OLM junction defects. Therefore, it remains possible that rescue of this underlying phenotype contributed to the improvements in photoreceptor survival and function that were observed in the *B-rescue* mice. Further work will be needed to evaluate the relative importance of the OLM and hemi-rosette pathologies. It is also possible that CRB1-B functions within the photoreceptor outer segment, which we previously found (3), and confirmed here, to be a site of CRB1-B protein expression. Aside from degenerative changes, we have not yet observed anatomical defects at the outer segment, but a role for CRB1-B at this site is yet to be ruled out. Finally, it was recently proposed that the key site of CRB1 pathology leading to degeneration is not in the eye, but rather the tight junctions of the gut (26). One limitation of the gut study is that it used *rd8* on the C57Bl6/N background as a disease model, which, as shown here and elsewhere (23), is a poor model for key disease features with minimal photoreceptor loss. Furthermore, the notion of gut pathology is hard to reconcile with our finding that expressing CRB1-B selectively in photoreceptors – but not in the gut (Supplemental Fig. 6E) – is sufficient to rescue degeneration.

### Therapeutic potential of CRB1-B

Even though the A and B isoforms are expressed in different cell types, loss-of-function mouse genetics suggest that removal of both isoforms is necessary for severe vision loss. Given these genetics, which imply that either the A or B isoform is sufficient to preserve visual function, it stands to reason that restoration of either isoform could be a viable therapeutic strategy. Nevertheless, there are several reasons to favor CRB1-B as the first-choice therapeutic candidate. First, it is the most abundantly expressed isoform in adult human retina, making it the obvious starting point for gene replacement therapy (3). Second, the *CRB1-B* open reading frame is short enough, at 3.0 kb, to fit into an AAV gene therapy vector, together with a cell type specific promoter and other elements promoting mRNA stability. *CRB1-A*, by contrast, is 4.2 kb, which leaves little room within the 4.7 kb AAV genome for these other key elements. One study found that AAV-mediated expression of CRB1-A was deleterious to photoreceptor survival in *ex1* mutant mice, suggesting that use of the A isoform in AAV therapeutics could be challenging (13). Here we show that transgenic restoration of CRB1-B expression by rod photoreceptors is sufficient to rescue visual acuity defects in *null* mutant mice. This finding suggests there is strong therapeutic potential in a rescue strategy centered on CRB1-B. Therefore, further work towards developing AAV-based rescue vectors could lead to a new gene therapy strategy for CRB1 disease.

## Methods

### Animals

Sex as a biological variable: Mice of both sexes were analyzed in this study and no systematic sex differences were found. Mice were subjected to a standard 12 h light-dark cycle with food and water access ad libitum. For all the experiments mice were anesthetized using isoflurane prior to decapitation.

C57Bl6/J and C57Bl6/N mice (carrying the *Crb1^rd8^* mutation) were purchased from Jackson Labs and Charles River respectively. The *Crb1^em1Jnk^*(i.e., *Crb1^ΔB^*) and *Crb1^em3Jnk^* (i.e., *Crb1^null^*) alleles, bearing CRISPR-Cas9 deletions of the genomic regions shown in Fig. 1C, were described previously (3). The *Crb1^ΔB^* allele eliminates exon 5a, the first exon of *Crb1-B*, while the *Crb1^null^* allele eliminates exons 5b, 6, and part of 7 including its splice acceptor site. The Rho-CRB1B transgenic line expresses mouse CRB1-B under control of the well characterized 4.4 kb mouse rhodopsin promoter fragment (27). Generation of this line is described below.

The *Crb1^ΔB^* and *Crb1^null^* mutant alleles, and the Rho-CRB1B transgenic line, were maintained by backcrossing to C57Bl6/J and are expected to be on a pure C57Bl6/J background since they were maintained this way for >10 generations (mutant alleles) or >5 generations (transgenic line) prior to use in this study. For loss-of-function genetic experiments, experimental mice of genotype n*ull/null*, *ΔB/ΔB*, and *null/ΔB* were generated by interbreeding *Crb1^null/+^* or *Crb1^ΔB/+^* heterozygotes; or in some cases by crossing these heterozygotes to *Crb1^null/null^* or *Crb1^ΔB/ΔB^* homozygotes. Strain-matched control mice were generated by first outcrossing *Crb1^null/+^*heterozygotes to C57Bl6/J, and then interbreeding the resulting *Crb1^+/+^*progeny.

For rescue experiments, Rho-CRB1B; *Crb1^null/+^* breeders were first generated by crossing the Rho-CRB1B and *Crb1^null^* strains. Breeders were generated in this manner in each generation, rather than by propagation through incrossing, to avoid genetic drift. To generate experimental animals, Rho-CRB1B; *Crb1^null/+^* breeders were subsequently crossed with *Crb1^null/null^* mice. One quarter of the progeny of this mating were *null* mutants without the transgene, and one quarter were *null* mutants carrying the transgene – i.e. *B-rescue* mice. To generate strain-matched wild-type control mice, the same Rho-CRB1B; *Crb1^null/+^*breeders were crossed to the *Crb1^+/+^* control strain, thereby generating *Crb1^+/+^* progeny with or without the transgene.

### Generation of CRB1-B rescue transgenic line

The DNA construct used for generating transgenic mice, pRho–Crb1-B–IRES–GFP, was generated as follows. First, a DNA fragment was synthesized (VectorBuilder) to contain three elements: 1) the mouse *Crb1-B* cDNA open reading frame; 2) an internal ribosome entry site (IRES) sequence; and 3) the *green fluorescent protein* open reading frame. This fragment was cloned into an expression vector (kind gift of J. Pearring and V. Arshavsky) containing the 4.4 kb rhodopsin promoter (20, 28). Linearized DNA was provided to the Duke Transgenic Core facility for generation of transgenic mice on the C57Bl6/J-SJL F1 hybrid background. Mice were backcrossed to C57Bl6/J and maintained on this background as noted above. Transgenic founders were identified by PCR using genotyping primers (Table 2) that flank the junction between the rhodopsin promoter and *Crb1-B* cDNA (expected band size 278 bp). Mice were subsequently genotyped using these same primers.

**Table 1.** Photoreceptor nuclei detection parameters.

|  |  |
| --- | --- |
| **Setup parameters** |  |
| Detection image | Optical density sum |
| Requested pixel size | 0.1625 $\mu\text{m}$ |
| **Nucleus parameters** |  |
| Background radius | 7 $\mu\text{m}$ |
| Median filter radius | 1 $\mu\text{m}$ |
| Sigma | 0.6 $\mu\text{m}$ |
| Minimum area | 1.5 $\mu\text{m}^2$ |
| Maximum area | 30 $\mu\text{m}^2$ |
| **Intensity parameters** |  |
| Threshold | 0.275 |
| Maximum background intensity | 1.6 |
| Split by shape | ✓ |
| **Cell parameters** |  |
| Cell expansion | 0 $\mu\text{m}$ |
| Include cell nucleus | ✓ |
| **General parameters** |  |
| Smooth boundaries | ✓ |
| Make measurements | ✓ |

Nucleus parameters are optimized for detecting all photoreceptors. These parameters are consistent between all images. The only parameter that can be changed for optimal detection is “threshold” because it is dependent on the intensity of the chromatin and variable staining between experiments can affect this parameter. Depending on whether the
nuclei are excluded or extra-nuclear material is detected, the “threshold” can either be increased or decreased step wise.

**Table 2.**
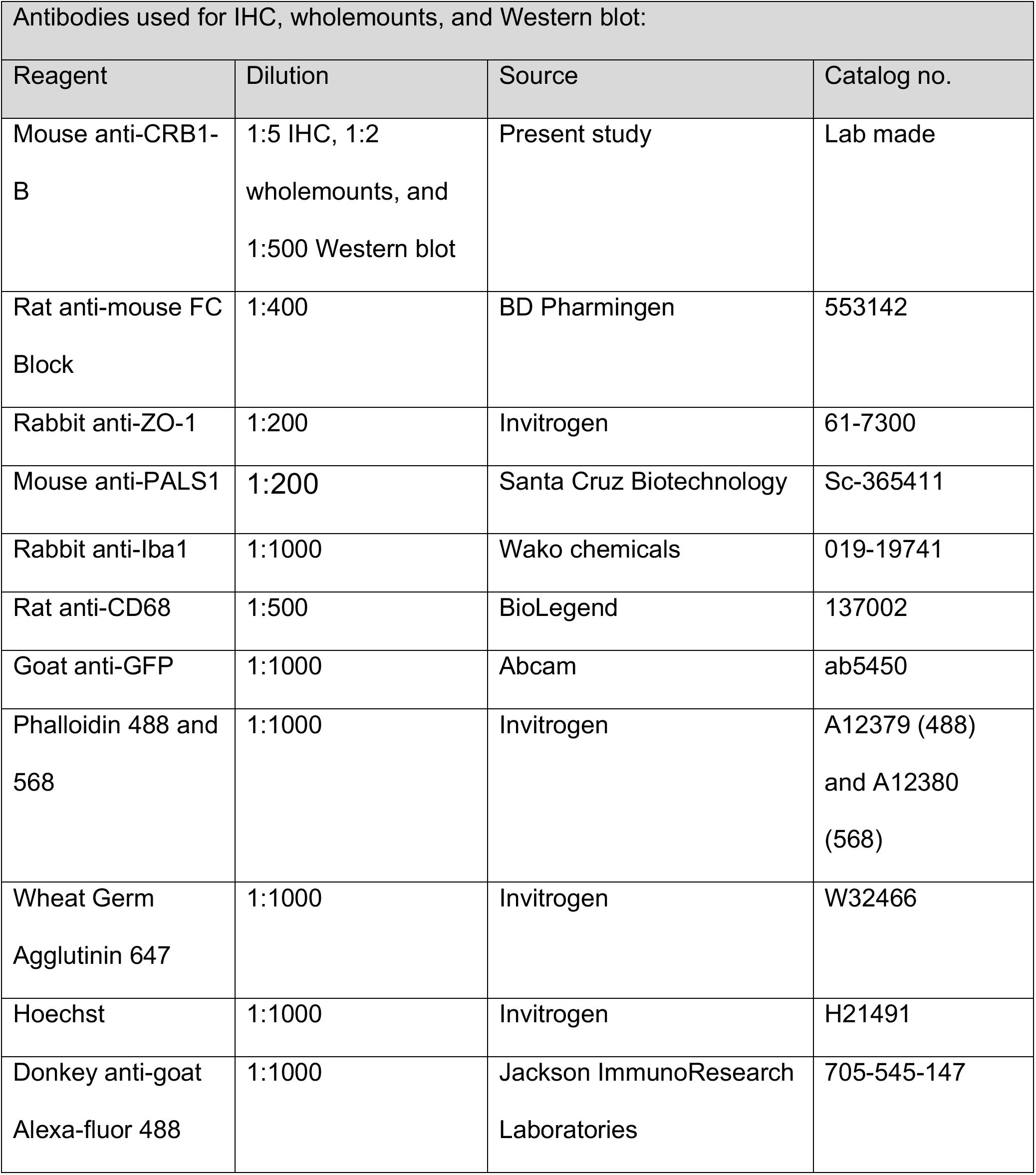
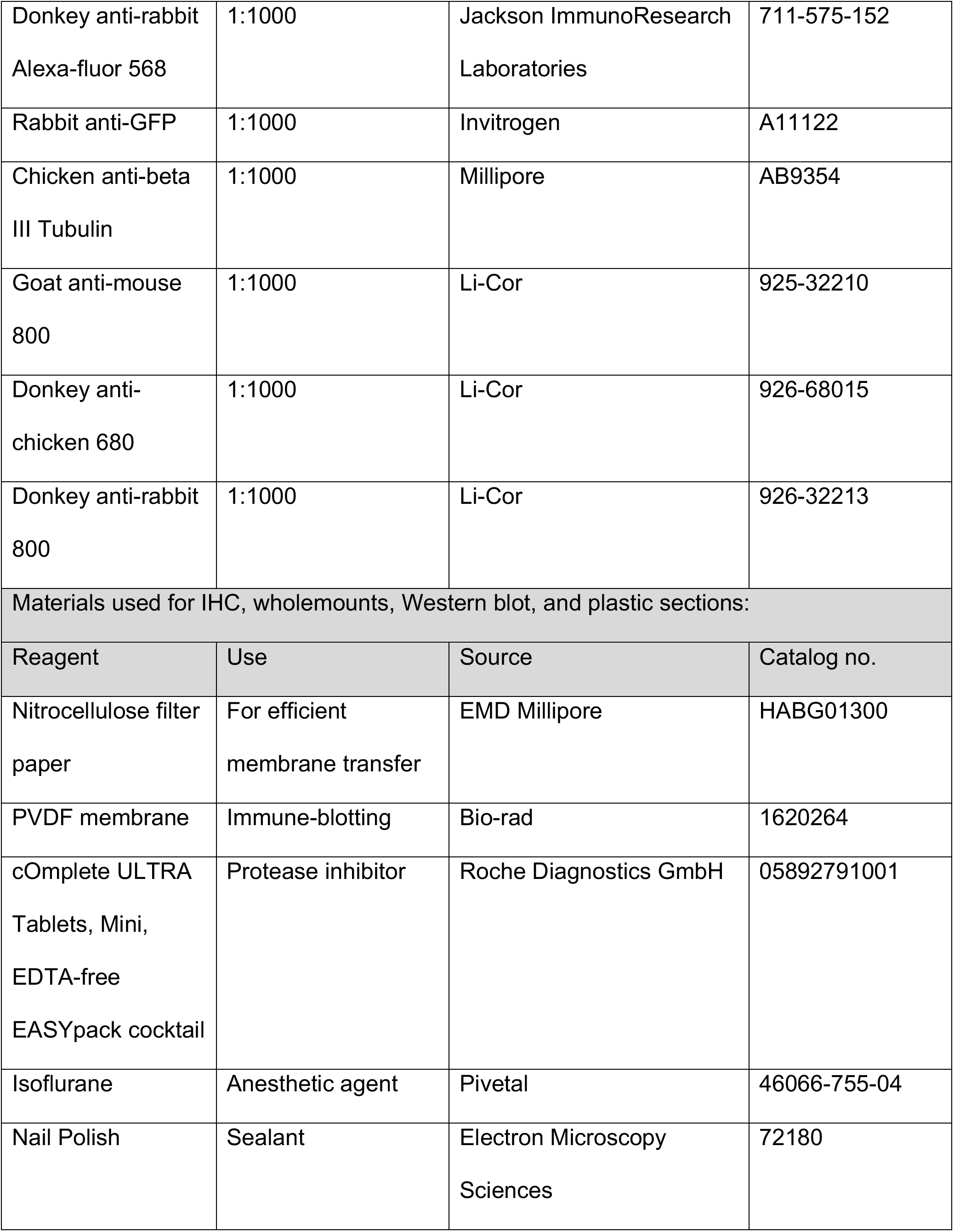
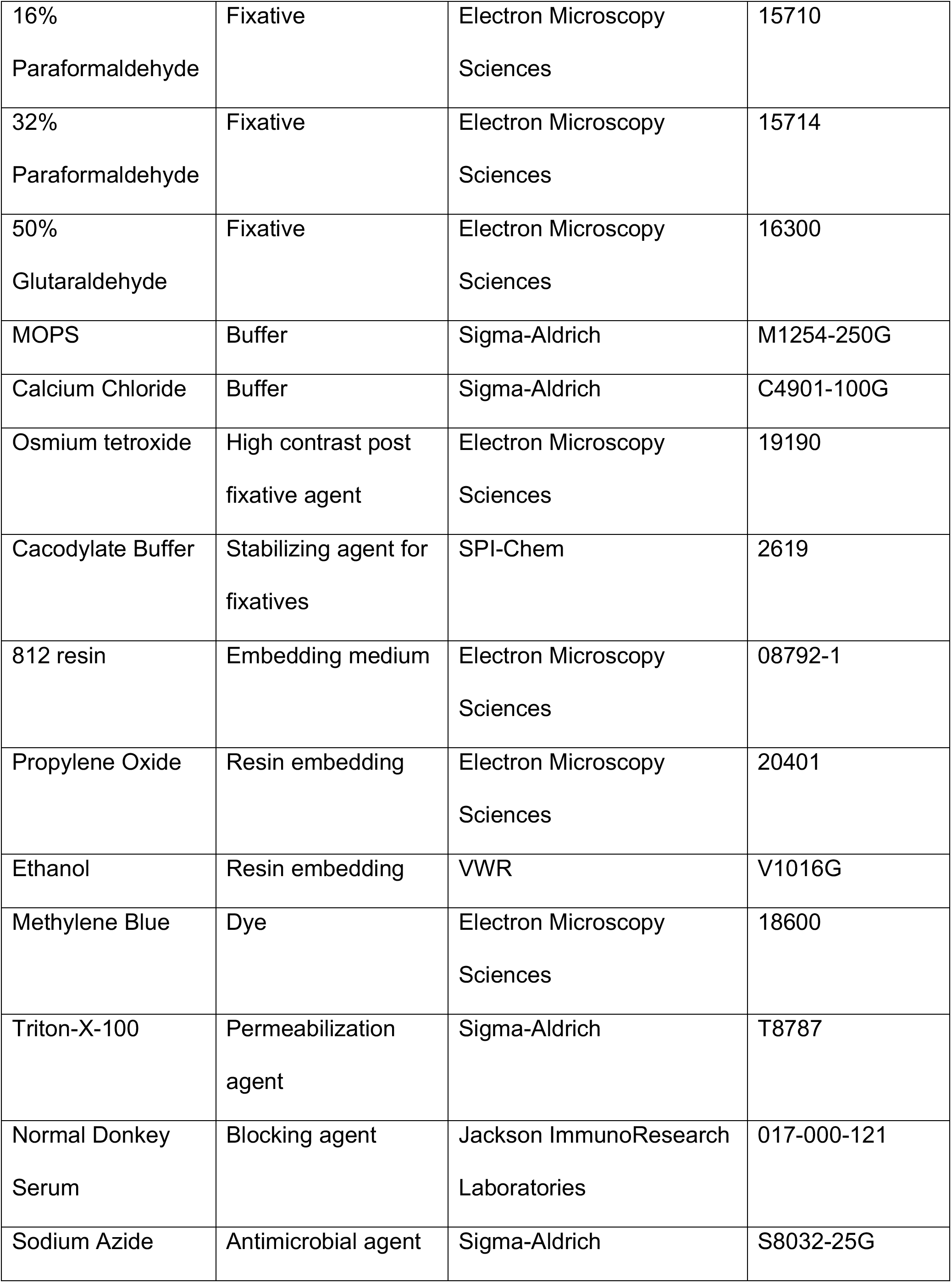
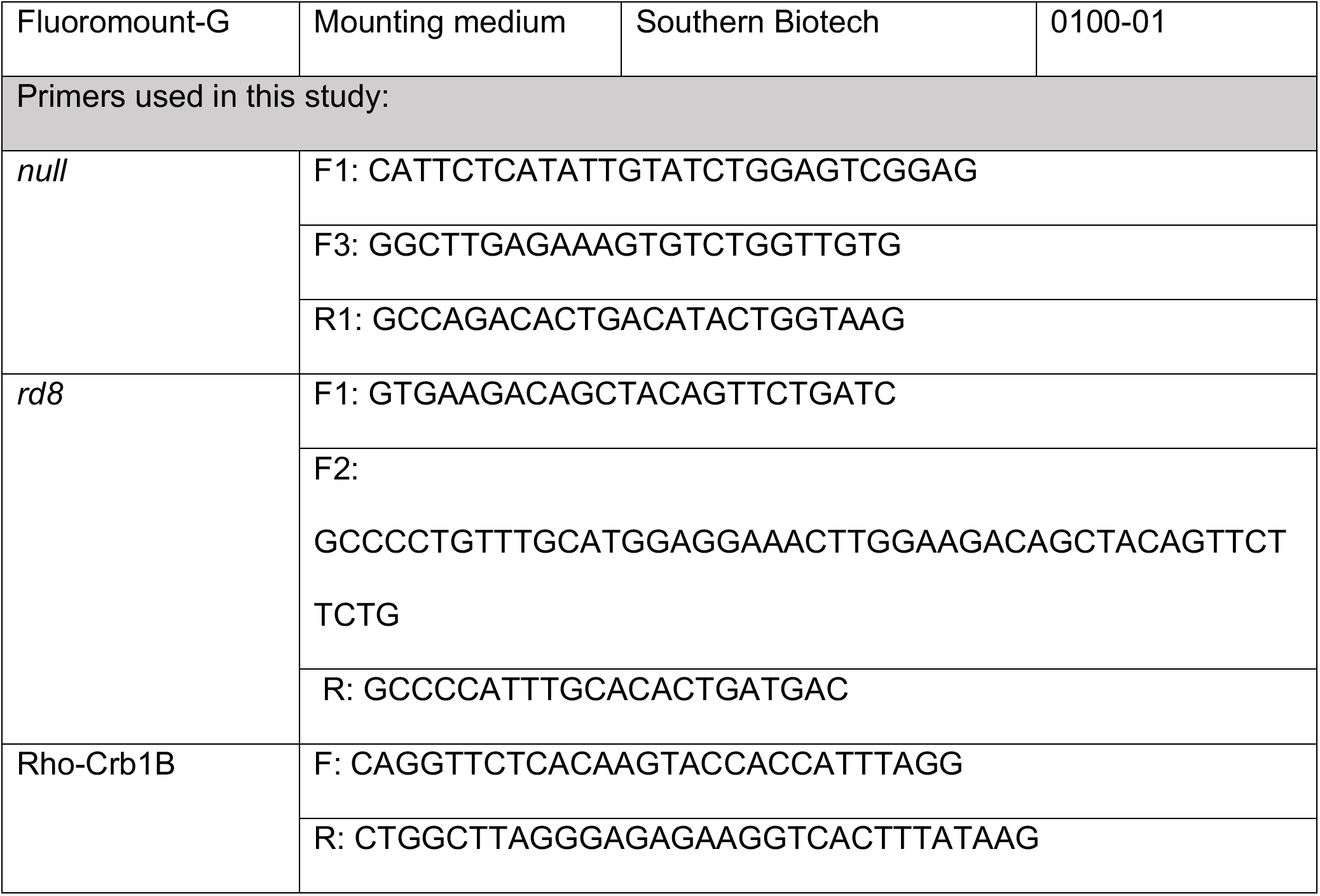
Key resources table.

### Crb1-B monoclonal antibody

A monoclonal antibody was generated by immunizing mice with a 16 amino acid peptide (RMNDEPVVEWGAQENY) comprising the entire intracellular domain of mouse CRB1-B (Absea Biotechnology Ltd., Beijing, China). Hybridoma clones were generated and screened by ELISA. Subsequently, to identify clones with high signal-to-noise ratios in various types of assays, supernatants from ELISA-positive clones (n =52) were obtained from Absea and tested by Western blotting of CRB1-B transfected HEK293T cells. Positive clones from this assay were further tested by Western blotting of mouse retinal lysates and immunohistochemistry on mouse retina vibratome sections, leading to a final selection of 13 high performing clones. These clones were then obtained from Absea and passaged multiple times in our lab to establish cell lines. Supernatants from clones 15D3, 4B8 and 9H7 were used for the studies here, including validation using *Crb1^null^* mice that the signals detected by Western blot and immunohistochemistry are specific (Fig. 3I, J; Fig. 6; Supplemental Fig. 2).

### Histology: Thin plastic sections

Eyes were collected from euthanized mice and marked at the superior region with a low-temperature cautery pen. Eyes were then fixed at 4°C overnight in fixation buffer (50mM MOPS pH 7.4, 2% glutaraldehyde, 2% paraformaldehyde, 0.05% CaCl2 in H2O), with rocking agitation. Following fixation, eyes were stored in PBS until further processing. For embedding, the eyecup was isolated and further fixed in a solution of 1% Osmium tetroxide in 0.1M cacodylate buffer. Eyecups were then dehydrated with graded Ethanol from 30% to 100% and propylene oxide, infiltrated overnight under the vacuum using a 1:1 mixture of propylene oxide and embed 812 resin, exchanged with embed 812 resin the next day, and finally were embedded in embed 812 resin at 65°C overnight. Semi-thin sections (0.5µm) were prepared using Leica Reichert Ultracuts, with sectioning oriented along the superior-inferior axis using the superior mark made at the time of tissue collection. Sections were stained with 1% Methylene Blue. Slides containing sections through central-most retina – containing the optic nerve head – were selected for downstream analysis.

### Photoreceptor counts

Rod and cone photoreceptors were counted in thin plastic sections prepared from central retina and oriented along the superior-inferior axis. 3 sections per eye were analyzed using the following pipeline, and averaged to obtain final rod and cone numbers for each eye.

Slides were tile scanned using a Leica Dmi8 inverted compound microscope equipped with a 40X/1.30 oil-immersion objective, using a LAS X acquisition software. Stitched images of full retinal sections were imported into Fiji/ImageJ (29). Regions of interest (ROIs) of 100µm width were placed along the OLM every 500 µm, with the first ROIs being 50µm inferior and superior to the optic nerve head. These ROIs were then imported into QuPath-0.5.1 software (30). Rod and cone nuclei were segmented by first annotating the image followed by running the following workflow: analyze → cell detection → cell detection → nuclei detection (parameters given in Table 1). These settings were established in pilot studies in which we optimized the parameters for detecting photoreceptor nuclei automatically. Once segmented and counted, nuclei were classified by hand as being from rods or cones, using established morphological criteria (see Supplemental Fig. 1A).

### Immunohistochemistry

For staining of vibratome retina cross-sections, eyes were fixed in 4% paraformaldehyde (PFA)/1x PBS for 1.5 hours on ice, washed in PBS twice and stored at 4°C in PBS + 0.02% sodium azide until further processing. Prior to sectioning, the anterior eye was removed together with the lens and vitreous humor. Posterior eyecup with retina was embedded in agarose at 60-65°C. From these embedded blocks, 100µm retinal sections were made using a Leica VT 1200 S vibrating blade microtome. Sections were blocked in PBS containing 0.3% Triton-X-100 and 3% normal donkey serum for 1 hour at room temperature on a rocker. Sections were then washed with 1X PBS+0.02% Sodium azide and incubated with antibodies to PALS1, GFP, CRB1-B and phalloidin diluted in blocking buffer overnight on a rocker at 4°C. Sections were then washed 3 times over a period of 30 min. and incubated with secondary antibodies diluted in 1X PBS for 1 hour at room temperature on a rocker. Sections were again washed 3-5 times over a period of 30 mins. with 1X PBS at room temperature on a rocker. Sections were gently placed on a microscope slide using a soft thin paint brush and coverslip was mounted using Fluoromount G.

For staining of retinal cryosections, eyes were fixed and stored as described above. After isolating the eyecup, it was submerged in 30% sucrose/PBS followed by embedding in Tissue Freezing Medium (VWR) and frozen in 2-methylbutane on dry ice. 20 µm sections were then cut using Leica CM 1950 cryostat and mounted on Superfrost Plus slides and further dried using a slide warmer. For immunolabeling, these slides were washed for 5 min with PBS and blocked using 1X PBS containing 0.3% Triton-X-100 and 3% normal donkey serum for 1 hour at room temperature. Antibodies to Iba1 and CD68 were diluted in blocking buffer. Slides were then incubated with these antibodies overnight at 4°C. Slides were washed with 1X PBS 2-5X for 10 min followed by incubation with secondary antibodies diluted in PBS + 0.3% Triton-X-100 for 1-2 hours at room temperature. Slides were washed 2-5X with 1X PBS for 10 min and then mounted using Fluoromount G.

For staining of retinal wholemounts for the phalloidin OLM assay, eyes were fixed as above and stored in PBS + 0.02% sodium azide until staining. The retina was dissected free of the eyecup and blocked for two hours on a rocker in PBS containing 3% normal donkey serum and 0.3% Triton-X-100. Retinas were then incubated in blocking buffer containing phalloidin and wheat-germ agglutinin (WGA); in some cases, primary antibodies to GFP or Iba1 were also included in the staining mixture. Staining proceeded for 5 days on a rocker at 4°C. Retinas were then washed 3-5 times over a period of 2 hours and incubated with secondary antibodies diluted in PBS + 0.3% Titon-X-100 at 4°C overnight on a rocker. Retinas were again washed 3-5 times with 1X PBS for 2 hours at RT on a rocker. Mounting was done by making 4 radial incisions in the retina, which was then flattened on a nitrocellulose filter paper with photoreceptor side facing up. The preparation was then coverslipped using Fluoromount G as the mounting media.

For staining of retinal wholemounts with antibodies to PALS1, ZO-1 and CRB1-B, the anterior portion of the enucleated eye was removed prior to fixation of the eyecup in 4% PFA/1x PBS for 30 min on ice. At the time of staining, the retina was separated from the eyecup and blocked for 30 min with 6% normal donkey serum + 0.5% Triton-X-100 in 1X PBS. Primary antibodies to PALS-1, ZO-1 and/or CRB1-B were diluted in PBS + 0.25% Triton-X-100 + 6% normal donkey serum + 0.02% sodium azide. Retinas were incubated in primary antibody solution for 2 days at 4°C on rocker. Retinas were then washed with 1X PBS 3-5 times at RT for 5 min each on a rocker. Secondary antibodies were incubated in the same solution as primary antibody for 2 days at 4°C on rocker. Retinas were again washed 3-5 times with 1X PBS at RT 5 minutes each on a rocker. Mounting of retinas was done in the same manner as above.

For all staining, Hoechst nuclear dye was included at the secondary antibody step as a counterstain. Samples were imaged using the Galvano scanner on a Nikon A1 confocal laser scanning microscope equipped with a motorized translational stage. Single slice snapshots and z-stacks were acquired using 60X oil objective (N.A. 1.49).

### Wholemount OLM assay

Phalloidin-stained retinal wholemounts were used to assess the extent of OLM damage in *Crb1* mutant mice. Labelled retinas were imaged on the confocal microscope as noted above at three eccentricities – center, middle, and periphery – as previously described (31). Briefly: Each retina was divided into 3 annular zones based on the distance between the optic nerve head and the retinal edge (i.e., the ONH-peripheral distance). The radius of each annulus was one-third of the full ONH-peripheral distance. ROIs for the central eccentricity were placed in the first annulus, while middle and peripheral ROIs were placed in the second and third zones. Four images were taken at each eccentricity in each retina, for a total of 12 ROIs per retina. Z-stacks with 0.5µm step size and X-Y dimensions of 212.13µm x 212.13 µm were acquired from each of these 12 locations.

To measure OLM holes, the Z-stacks were analyzed in Fiji/ImageJ. The number of holes was counted over the entire Z-stack manually using the Fiji multipoint tool. Holes that contacted one side of the field of view were excluded from the analysis.

To measure the size of individual holes, we examined the subset of Z-slices encompassing the OLM (usually 4-5 Z planes) and marked each individual hole within one of these slices using the multipoint tool. Holes were then segmented using the Fiji Level Sets tool. Because a whole-mounted retina is never perfectly flat, some holes appeared in multiple Z slices. In these cases, a single slice was chosen to represent the hole, to avoid double measurement. Graphical representations of hole burden (e.g. Fig. 3D) were generated by Z-projection of the segmented holes across all slices. Areas of the segmented holes were used to generate hole size histograms on a per-animal basis. The cumulative histograms shown in the figures give the mean ± S.E.M. for each hole size bin across all animals.

To compute OLM hole burden (i.e., the fraction of OLM area occupied by holes), the areas of all holes within an individual field of view were summed and divided by the field of view area. This procedure yielded 12 separate measurements of OLM hole burden for each animal, which were averaged – either per eccentricity or across all 12 images – to yield the fractional hole coverage reported for each animal.

### Visual behavior using OptoMotry

Mice were dark-adapted for at least 2 hours prior to starting the procedure. Visual threshold/visual acuity were measured by a modified OptoMotry system (Cerebral Mechanics, Medicine Hat, Alberta, Canada) for opto-kinetic tracking in a rodent as described previously (32). Reduced luminance was set by layering combinations of neutral density (ND) filters (Lee Filters USA, Andover, Hampshire, UK) on the motor screens. Two infrared lights were installed inside the chamber, to visualize the mice movement. All the tests performed involved a default program with fixed speed (12 degrees/second) and mesopic condition (4.2 ND, about 1.7X10^−3^ cd/m^2^) as measured by a light meter (ILT-1700; International Light Technologies, Peabody, MA, USA). The screens generate a virtual cylinder comprised of a vertical sine wave grating that rotates at a constant speed surrounding the animal. Mice were allowed to move freely on the platform and a video camera fixed above the animal allowed the investigator to determine if the animal made a tracking motion with reflexive head and neck movements in response to the visual stimuli. Spatial frequency threshold was quantified by increasing the spatial frequency of the grating until the animal no longer responded. The maximum spatial frequency that still generated a tracking response provided the measure of visual acuity. Animals were allowed to acclimate to the testing apparatus during a mock testing session which was conducted two weeks prior to the actual testing session. Acuity measurements were collected separately for stripes moving in both directions, but the data were averaged to give the final acuity measurement for each animal.

### Western blotting

Retinas and gut from Crb1 wild-type, *null*, and *B-rescue* mice, aged approximately P30, were dissected out in ice-cold Ringer’s solution. Retinas were lysed in 1% DM solution containing 0.1% w/v dodecyl maltoside, 50mM HEPES, 140mM NaCl, 3mM MgCl2, and 1 cOmplete^TM^ ULTRA Tablet, Mini, EDTA-free, EASYpack Protease Inhibitor Cocktail. Equal volumes of the lysates from respective genotypes were subjected to SDS-PAGE and proteins were transferred to polyvinylidene fluoride membranes. The membranes were blocked in the Li-Cor blocking buffer and incubated with the anti-CRB1-B, anti-GFP, and anti-ß-Tubulin antibodies overnight at 4°C. Subsequently, membranes were incubated with secondary antibodies IRDye®800CW goat anti-mouse, IRDye®680RD donkey anti-chicken, IRDye®800CW donkey anti-rabbit and imaged with Li-COR Odyssey CLx system.

### Statistics

All statistical analysis were performed using GraphPad Prism software (version11.0.0). *P* value <0.05 was considered significant. Group means were compared using either one-way ANOVA, two-way ANOVA, unpaired *t* test, Mann-Whitney test or Kolmogorov-Smirnov test. Details of specific statistical test are mentioned in the respective figure legends.

### Study approval

Animal work was reviewed and approved by the Duke University Institutional Animal Care and Use Committee.

## Data availability

Data underlying all the figures is submitted in a separate excel file with corresponding figure tabs.

## Author contributions

Conceptualization: E.M.D. and J.N.K.; methodology: E.M.D., C.K., M.D.; experiments and data analysis: E.M.D., J.V.L., C.K., L.Y., C.R., M.D., D.S., Y.H. and A.P.; writing-original draft: E.M.D. and J.N.K.; writing-review and editing: E.M.D., J.V.L., M.D., C.K., and J.N.K.; visualization: E.M.D. and J.N.K.; supervision: J.N.K.; project administration: J.N.K.; funding administration: E.M.D. and J.N.K.

## Funding support

This work was supported by the National Eye Institute (EY035637 to J.N.K.; EY035119 to J.C.V.-L.; EY5722 to Duke University), by Foundation Fighting Blindness (grants BR-CMMM-0619-0767-DUKE and PPA-1224-0890-DUKE to J.N.K.), by Research to Prevent Blindness (Stein Innovation Award to J.N.K. and an unrestricted grant to Duke University); and by Ruth K. Broad Biomedical Research Foundation (Postdoctoral Fellowship award to E.M.D.).

## Acknowledgements

We would like to thank Duke University Microscopy Core for providing access to the imaging facility and Duke University Transgenic and Knockout Mouse Core for assistance with generation of transgenic mouse. We thank Nicholas Schneider for providing expertise with vibratome sectioning. We thank Vadim Arshavsky. Oleg Alekseev, and all Kay Lab members for their helpful comments on the manuscript.

## Declaration of competing interests

JNK is a consultant for Retinaria, Inc. and is an inventor on patent US20220125948A1. All other authors declare no competing interests.

## Notes

### Summary of Updates

Figure changes: Model figures moved to a new Figure 1; layout and organization of several figures updated to improve clarity and narrative flow. Supplemental figures updated. Minor changes to the manuscript to fix typos and enhance clarity. Funding sources updated to fix inadvertent omission from original submission.

