## Supplemental Figures and Legends for "Retinal degeneration in a mouse model of CRB1 disease rescued by the photoreceptor-specific *CRB1-B* isoform"

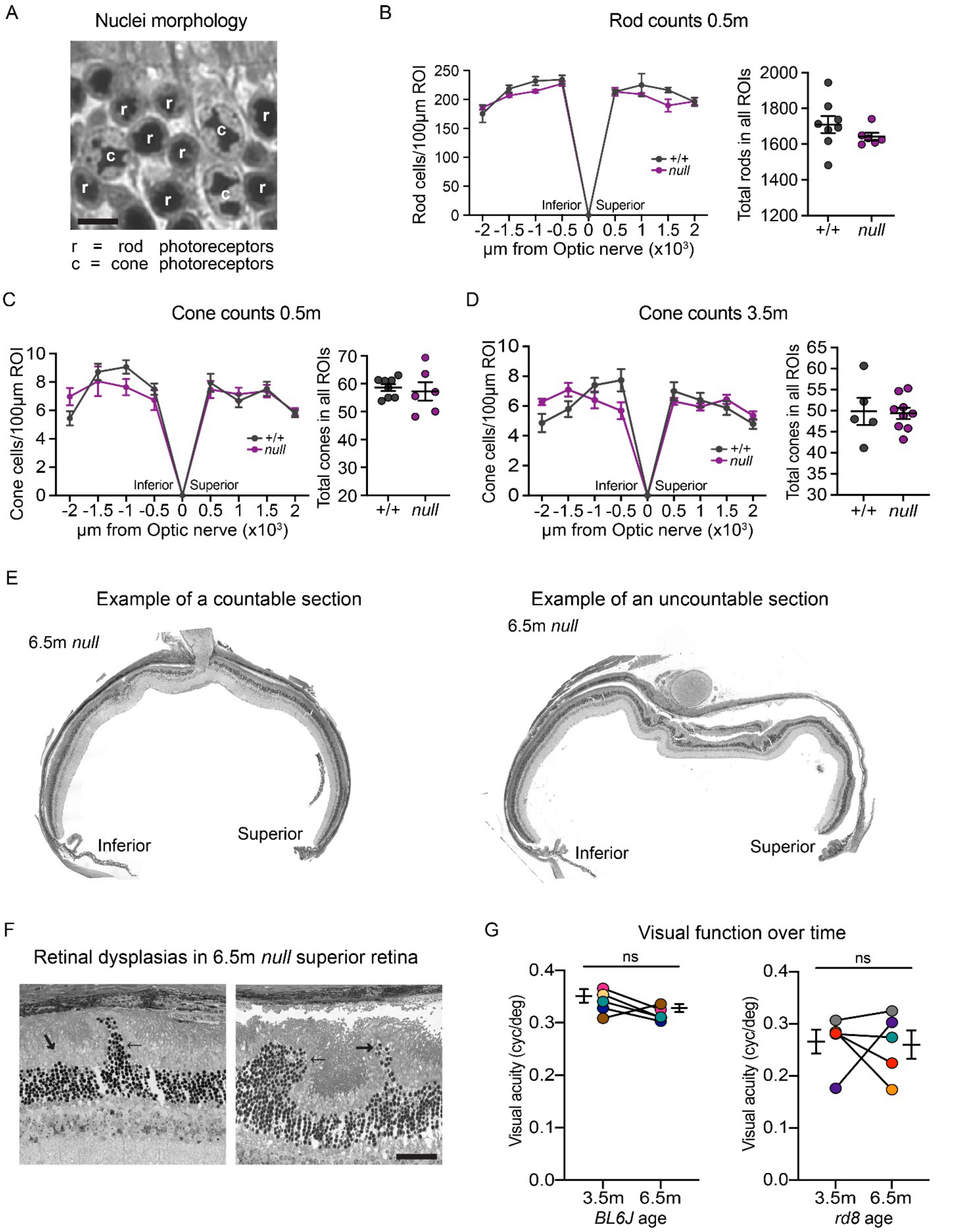


**Supplemental. Fig. 1:** Anatomical and behavioral analysis supporting Figure 2

(A) Identification of rods and cones for nuclei count analysis. Rods were identified by their characteristic round nuclei and compact chromatin pattern, and cones were identified by their comparatively large oval/conical nuclei and the segmented/irregular chromatin arrangement.

(B) Rod photoreceptor cell counts at 0.5m. Left, spider plot; right, total cones counted across all ROIs. Statistics: (A) Spider plot: (left) two-way Anova (ns $p = 0.069$); (right) Total rod count: unpaired $t$ test, ns $p = 0.272$.

(C) Cone photoreceptor cell counts at 0.5m. Left, spider plot; right, total cones counted across all ROIs. Statistics: (A) Spider plot: (left) two-way Anova (ns $p = 0.541$); (right) Total cone count: unpaired $t$ test, ns $p = 0.676$.

(D) Cone photoreceptor cell counts at 3.5m. Left, spider plot; right, total cones counted across all ROIs. Statistics: (A) Spider plot: (left) two-way Anova (ns $p = 0.851$); (right) Total cone count: unpaired $t$ test, ns $p = 0.878$.

(E) Thin plastic sections of 6.5m Crb1 *null* retinas illustrating phenotypes that prevented ONL counts. Left, typical *null* phenotype. Right, an example of a retina that was excluded from the analysis. The counting strategy assumes that ROIs represent cross-sections perpendicular to the plane of the ONL. The large-scale folding and warping of the ONL evident in the excluded retina break this assumption and necessitated its exclusion from the cell counts. However, this phenotype could still reflect a severe impact on photoreceptor survival and function, leading us to conclude that cell counts (Fig. 2E,F) may have underestimated the severity of the 6.5m phenotype.

(F) Higher-magnification views of large retinal dysplasias (arrows) found in Crb1 *null* superior retina at 6.5m. ROIs for cell counting were placed to avoid these regions. The geometry of these dysplasias, and their presence in superior retina, distinguishes them from the hemi-rosettes previously described for other alleles in inferior retina (Mehalow et al., 2003; van de Pavert et al., 2004), which were also observed at 3.5m in *null* mice (Fig. 2G).

(G) Optokinetic response of BL6J (left, replotted from Fig. 2C) and *rd8* (right) at 3.5m vs 6.5m. Neither strain shows significant decline in visual function between these time points. Statistics: paired $t$ test, $p$ ns (BL6J) = 0.159, ns (*rd8*) = 0.887.

Error bars: mean ± S.E.M. Sample sizes: (B, C) n = 8 +/+, 6 *null* (D) n = 5 +/+, 9 *null*; (G) n = 5 BL6J, 5 *rd8*.

Scale bar: (B) 5 µm; (D) 50 µm.

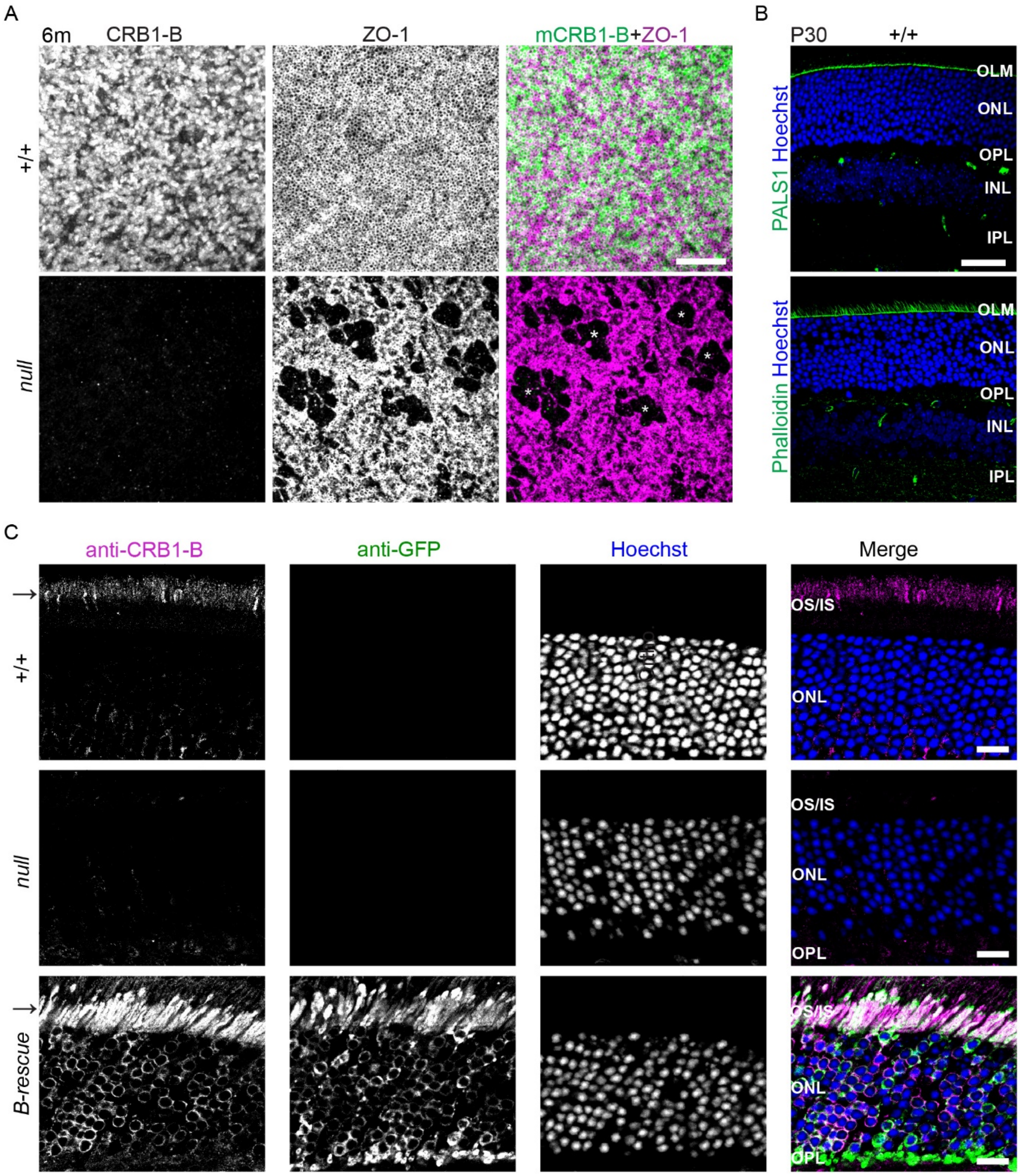


**<u>Supplemental Fig. 2</u>**<u>: Characterization of anti-CRB1-B monoclonal antibody and other OLM markers</u>

(A) *En face* confocal images showing the OLM in 6m Crb1 +/+ (top) and *null* (bottom) wholemount retina. Each panel is a maximum-intensity projection prepared from a Z-stack encompassing the OLM; only the slices at the OLM were projected. OLM is labelled using ZO-1 and co-labelled with mCrb1-B lab-made antibody. Crb1 +/+ retina shows a net-like pattern of ZO-1 and presence of Crb1-B in IS/OS as well as at OLM which is a characteristic of a healthy OLM. On the contrary, in *null* retina the OLM shows missing adherens junctions at multiple cites identified by ∗ with a complete absence of Crb1-B in IS/OS as well as OLM.

(B) Vibratome section images acquired at confocal microscope from P30 Crb1 +/+ retina showing the intact retinal architecture. Top image shows the OLM labelled by a known OLM marker, PALS1 (green). Bottom image shows the OLM labelled by Phalloidin (green). Both PALS1 and Phalloidin label the OLM supporting our experimental paradigm in this study for using Phalloidin as an OLM marker.

(C) Representative images of vibratome sections from Crb1 +/+ (top), *null* (middle), and *B-resc* (bottom) retinas, labelled with lab-made anti-CRB1-B antibody (magenta) and co-labelled with anti-GFP (green). Note the immunolabelling of Crb1-B (→) is only visible in Crb1 +/+ and *resc* retina. Crb1-B immunosignals are absent from the *null* retina. GFP immunosignals in the *resc* retina shows an adequate expression of the rescue transgene.

Scale bars: (A) 20 µm; (B, C) 30 µm.

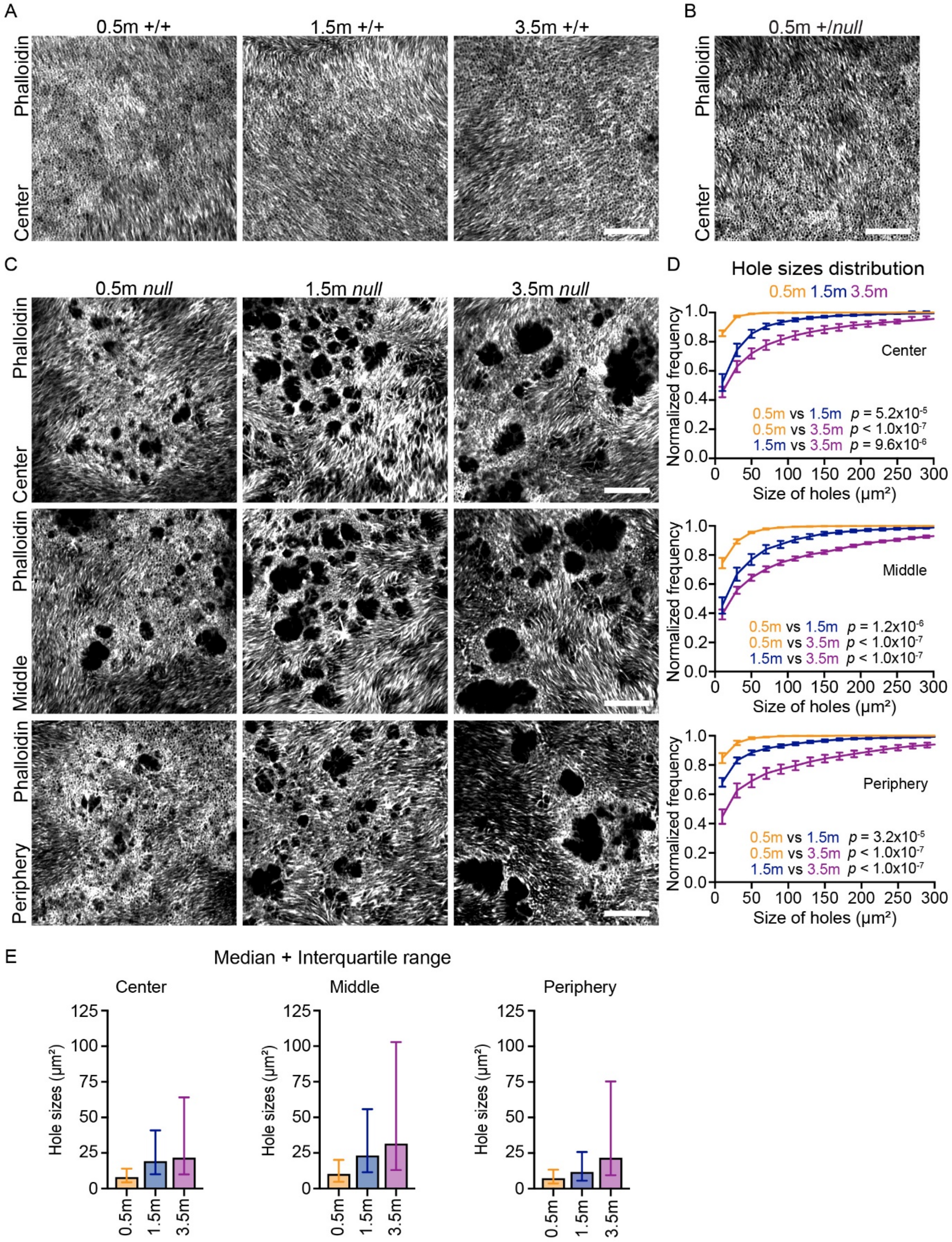


**Supplemental Fig. 3**: OLM anatomy in *Crb1*$^{null}$ and control mice

(A, B) Crb1 control mice lack OLM holes. Confocal images showing *en face* views of central retina OLM in +/+ mice (A), or *null/*+ heterozygous mice (B), at the indicated ages.

(C, D) Analysis of *Crb1*$^{null}$ OLM hole sizes over time at different retinal eccentricities. C: Confocal images of phalloidin-stained wholemount retinas showing representative en -face views of *null* mutant OLM at the indicated ages and the indicated retinal locations (see Methods for definitions of Center, Middle, and Peripheral). D: Quantification of hole sizes from images similar to C. Cumulative distribution histograms (truncated at 300 μm) depict changes in hole sizes over time. Peripheral retina is initially less affected at young ages. However, holes become progressively larger over time at each eccentricity. Error bars: mean ± S.E.M. Statistics: Kolmogorov-Smirnov test for frequency distribution. Top to bottom: Center, Middle and Periphery with *p* values for all comparisons shown on graphs. (E) Distribution of hole sizes (untruncated) represented as median (column bars) and interquartile range (IQR, whiskers – lower $25^{th}$ and higher $75^{th}$ percentile) in at indicated ages and retinal location.

Sample size: *null* 0.5m n = 5; 1.5m n = 7; 3.5m n = 7.

Scale bars: (A-C) 20 μm.

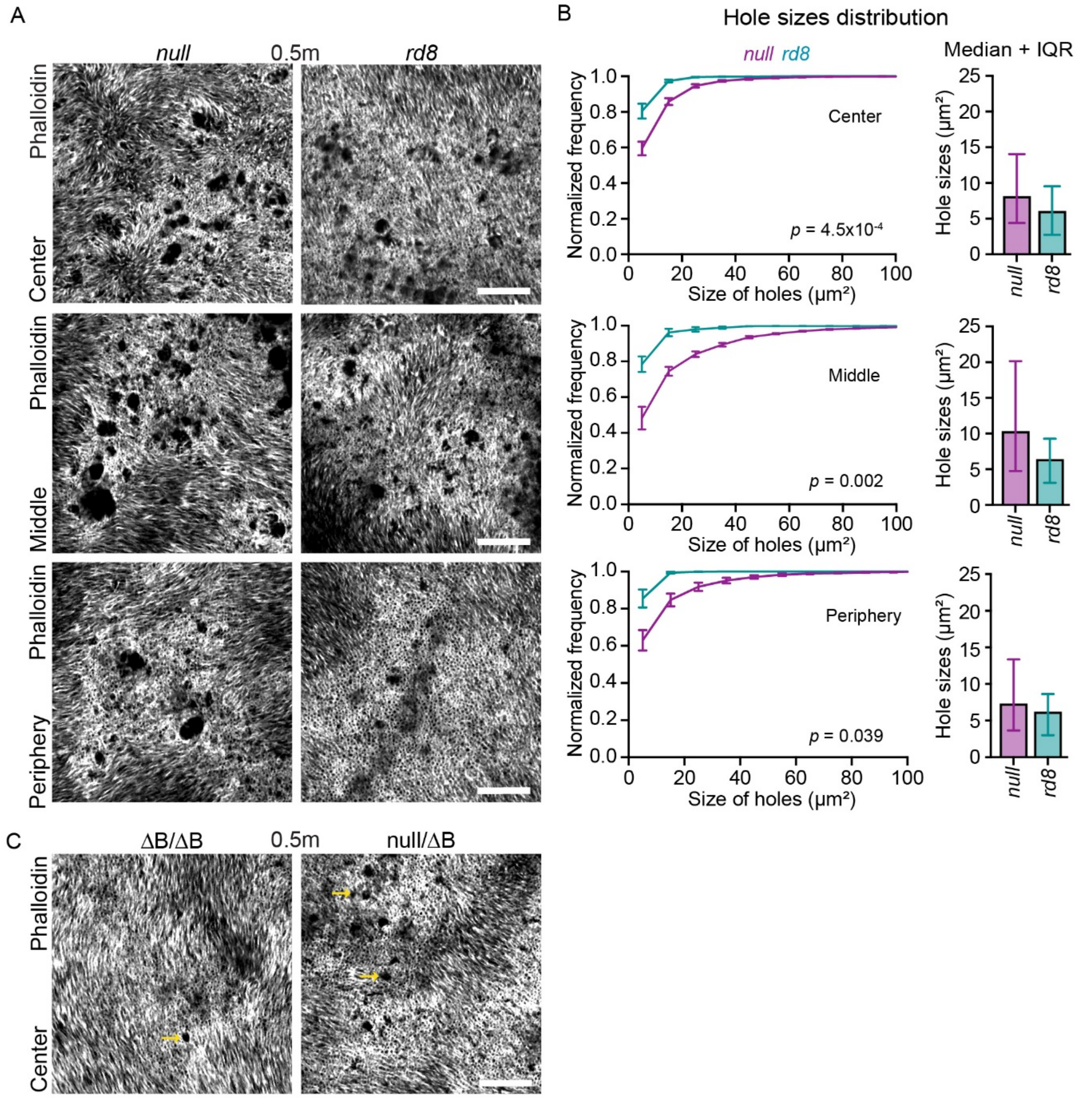


**Supplemental. Fig. 4:** OLM phenotypes in Crb1 *null*, *rd8* and ΔB mutant alleles

(A) Confocal images showing en -face view of OLM in 0.5m Crb1 *null* mice (left) and *rd8* mice (right). Images were taken from central retina (top), middle, and peripheral (bottom) retinal eccentricities.

(B) Left, cumulative distribution histograms of holes sizes (truncated at 100 µm) at each eccentricity, quantified from images similar to A. OLM of *null* mutants (magenta) have significantly larger holes than *rd8* (teal) at all eccentricities, although the difference is biggest in central retina (top). Error bars: mean ± S.E.M. Right, hole sizes distribution (untruncated) represented as median (column bars), and interquartile range (IQR, whiskers – lower 25$^{th}$ and higher 75$^{th}$ percentile). Statistics: Kolmogorov-Smirnov test for frequency distribution. Top to bottom: Center, Middle and Periphery with *p* values for all comparisons shown on graphs.

(C) Representative en face views of OLM holes in 0.5m Crb1 ΔB/ΔB and Crb1 null/ΔB mice. Arrows points to characteristic small OLM holes in both genotypes.

Sample size: n = 5 *null*; n = 5 *rd8*. Scale bars: (A, C) 20 µm.

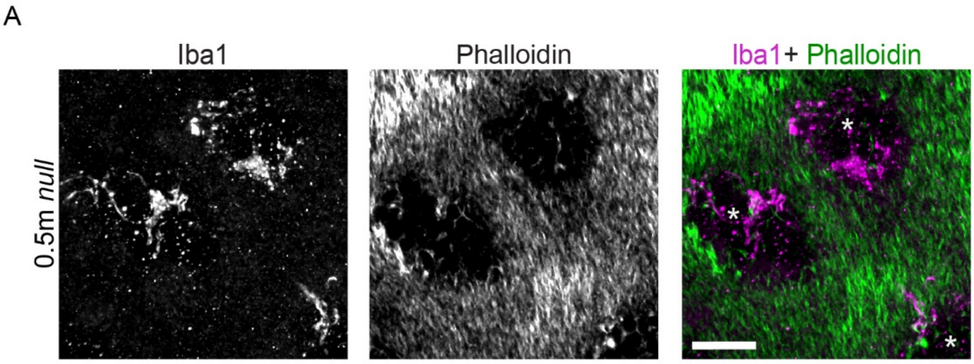


**Supplemental Fig. 5:** Microglia at OLM holes in young *Crb1*$^{null}$ mice

*En-face* confocal image of OLM from a 0.5m *null* wholemount retina, labelled with Phalloidin to mark OLM junctions and anti-Iba1 to mark microglia. Note that the microglia arbors ramify within areas of missing junctions (asterisks). This 0.5m mutant was more severely affected by OLM holes than the typical *null* animal at this age (see Fig. 3). We include it here to show that OLM pathology can start as early as 2-3 weeks of age in some *Crb1* mutant mice. Scale bar: 20 μm.

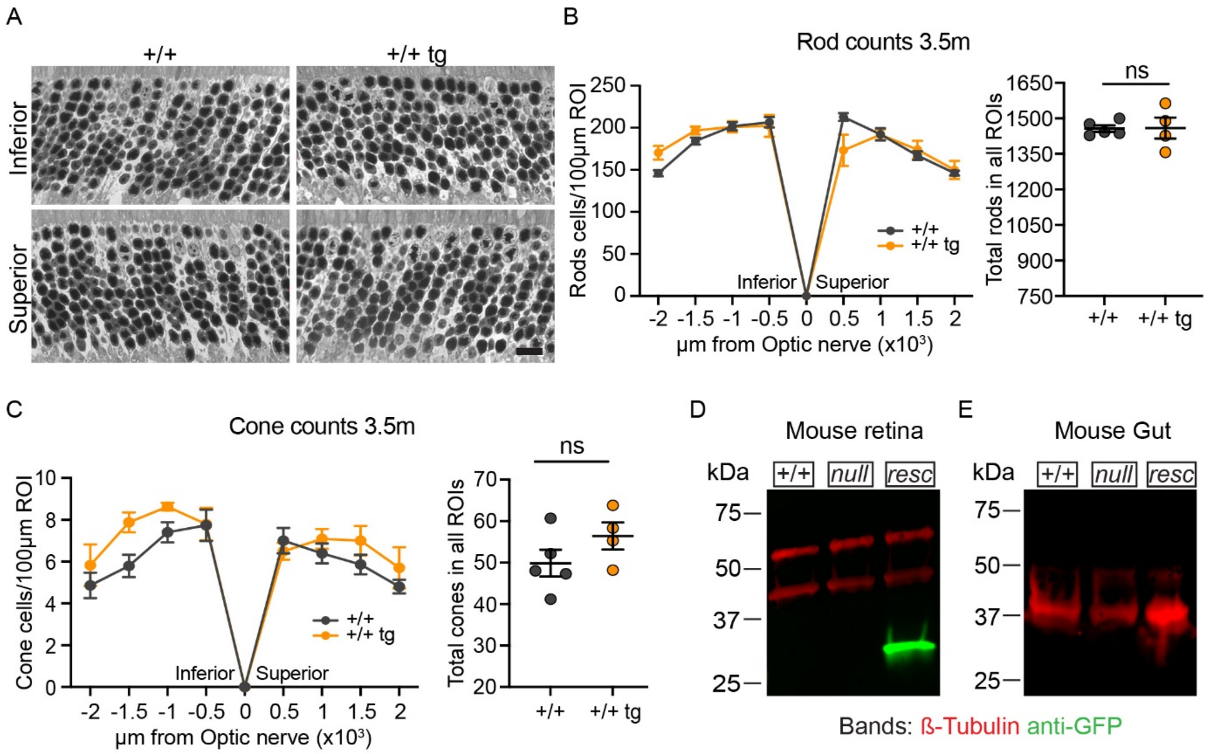


**<u>Supplemental Fig. 6:</u>** <u>Characterization of Rho-CRB1B-IRES-GFP rescue transgene</u>

(A, B) Rho-CRB1B transgene is not toxic to rod photoreceptors. A, Representative thin plastic section images of ONL from 3.5m Crb1 +/+ mice carrying Rho-CRB1B transgene (+/+ Tg, right) and non-transgenic +/+ controls (left). B, Quantification of rod nuclei from the images similar to A, acquired at 500µm intervals across the inferior-superior axis. Left, spider plot; right, total rod nuclei summed across all ROIs. Images in A are representative of the ROIs acquired from the two central region bins in B closest to the optic nerve head (Inferior, -500µm; superior, +500µm). Statistics: Spider plot (left): +/+ vs +/+ tg, two-way ANOVA, ns $p$ = 0.944. Total rod count (right): Mann-Whitney test, ns $p$ = 0.905.

(C) Rho-CRB1B transgene is also not toxic to cones. Left, spider plots showing cone photoreceptor counts in the same series of images used for rod counts (A, B). Right, total cones summed across all ROIs. Statistics: Total cone count (right): Mann-Whitney test, ns $p$ = 0.190.

(D, E) Western Blot analysis using anti-GFP to confirm retinal transgene expression. Lysates from retina (D) or gut (E) of Crb1 +/+, *null*, and *B-rescue* (resc) mice were probed using anti-GFP and anti-ß-Tubulin. The *resc* retina sample shows GFP expression from the *IRES-GFP* portion of the transgene at the expected size (~27 kDa) whereas non-transgenic +/+ and *null* retinas lack the GFP expression (D). GFP was not detected in gut. ß-Tubulin bands show equal loading of protein in all lanes.

Error bars: mean ± S.E.M.

Sample size: n = 5 +/+, n = 4 +/+ *B-rescue*.

Scale bar: (A) 10 µm; (D) 30 µm.

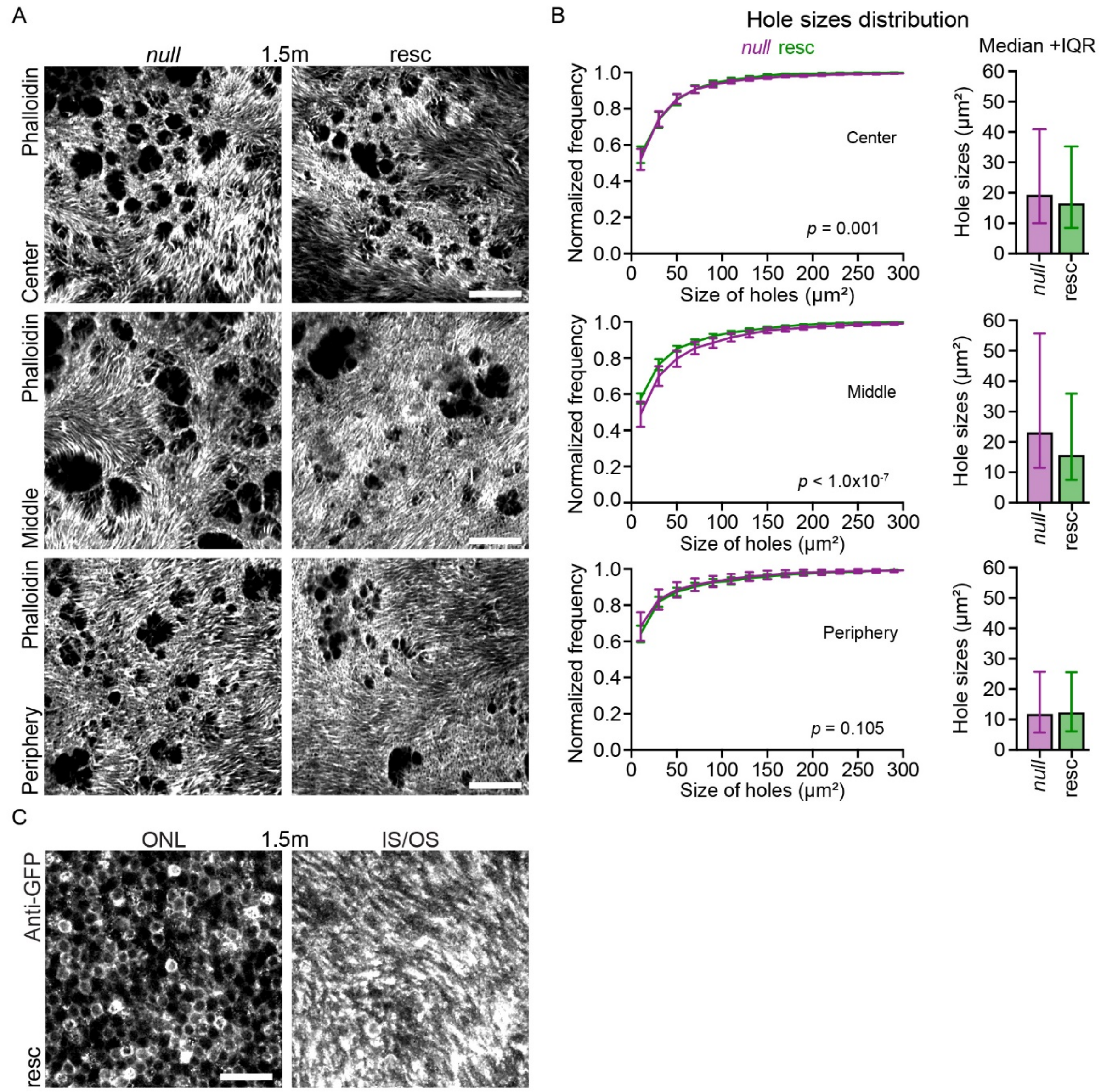


**Supplemental Fig. 7:** OLM hole phenotypes in *null* and *B-rescue* mice at 1.5m.

(A) En face view of OLM at central, middle, and peripheral eccentricities in 1.5m Crb1 *null* and *B-rescue* mice. Images are single confocal slices acquired from wholemount retinas labelled by Phalloidin.

(B) Left, cumulative distribution histograms (truncated at 300 µm) of hole sizes for each eccentricity quantified from image similar to A, comparing Crb1 *null* (magenta) to *B-rescue* mice (green). Error bars: mean ± S.E.M. Right, hole sizes distribution (untruncated) represented as median (column bars), and interquartile range (IQR, whiskers – lower 25$^{th}$ and higher 75$^{th}$ percentile). Statistics: Kolmogorov-Smirnov test for frequency distribution. Top to bottom: Center – ∗∗; Middle – ∗∗∗∗; Periphery – ns*;* for *p* values see graph.

(C) Confocal images of a representative B-recue retinal wholemount confirming transgene expression by photoreceptors. GFP labeling is visible in rods at the nuclear (ONL) and inner/outer segment (IS/OS) levels.

Sample size: n = 7 *null*; n = 7 *B-rescue*. Scale bars: (A, C) 20 µm.

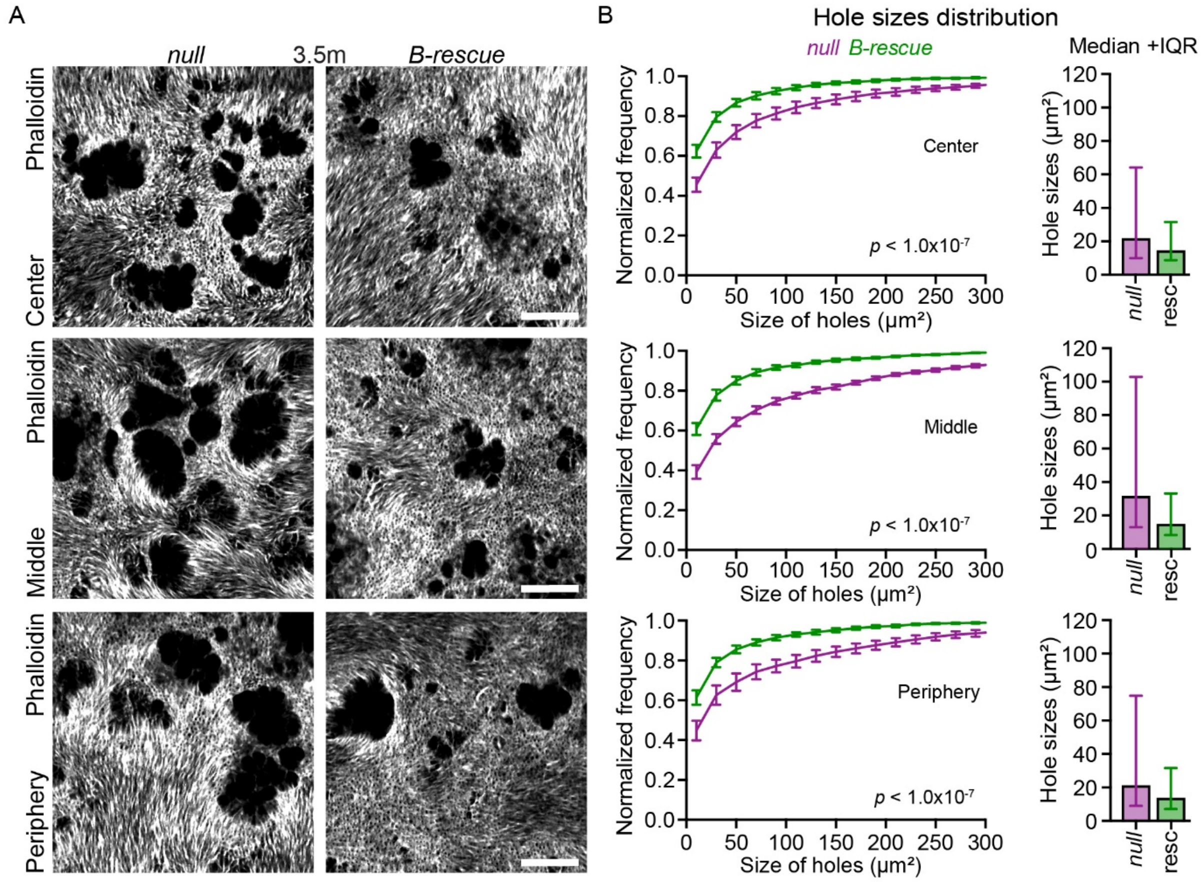


**Supplemental Fig. 8:** OLM hole phenotypes in *null* and *B-rescue* mice at 3.5m.

(A) En face views of OLM central, middle, and peripheral eccentricities in 3.5m Crb1 *null* and *B-rescue* mice. Images are single confocal slices acquired from wholemount retinas labelled by Phalloidin.

(B) Left, cumulative distribution histograms (truncated at 300 µm) of hole sizes for each eccentricity, quantified from images similar to A, comparing Crb1 *null* (magenta) to *B-rescue* (green). Error bars: mean ± S.E.M. Right, hole sizes distribution (untruncated) represented as median (column bars), and interquartile range (IQR, whiskers – lower $25^{th}$ and higher $75^{th}$ percentile). Statistics: Kolmogorov-Smirnov test for frequency distribution. Top to bottom: ∗∗∗∗ *p* values for all comparisons shown on the graph.

Data: mean ± S.E.M.

Sample size: n = 7 *null*; n = 7 *B-rescue*.

Scale bar: (A) 20 µm.
